# The neuropeptide SMYamide, a SIFamide paralog, is expressed by salivary gland innervating neurons in the American cockroach and likely functions as a hormone

**DOI:** 10.1101/2020.09.17.302331

**Authors:** Jan A. Veenstra

**Affiliations:** INCIA UMR 5287 CNRS, Université de Bordeaux, Pessac, France

**Keywords:** SIFamide, SMYamide, salivary gland, innervation, hormone, gonad

## Abstract

The SMYamide genes are paralogs of the SIFamide genes and code for neuropeptides that are structurally similar to SIFamide. In the American cockroach, *Periplanea americana*, the SMYamide gene is specifically expressed in the SN2 neurons that innervate the salivary glands and are known to produce action potentials during feeding. The innervation of the salivary glands by the SN2 neurons is such that one has to expect that on activation of these neurons significant amounts of SMYamide will be released into the hemolymph, thus suggesting that SMYamide also functions as a hormone. In the *Periplaneta* genome there are two putative SIFamide receptors and these are both expressed not only in the central nervous system and the salivary gland, but also in the gonads and other peripheral tissues. This reinforces the hypothesis that SMYamide also has an endocrine function in this species.

## 1. Introduction

SIFamide is an arthropod neuropeptide that was initially identified from a flesh fly by its ability to stimulate oviduct contractions in a locust. Antiserum raised to this neuropeptide showed it to be present in four large cells in the *pars intermedia*, a brain nucleus containing various types of neuroendocrine cells that project to the corpus cardiacum where hormones are released into the hemolymph. However, no SIFamide immunoreactivity was found in the corpus cardiacum (Janssen et al., 1996). We isolated and sequenced the peptide from the fruit fly *Drosophila melanogaster*. As in the fleshfly only four large cells in the brain were found to produce the peptide as shown by immunohistology, *in situ* hybridisation and transgene expression (Terhzaz et al., 2007). In the larva these neurons project a major axon to the ventral neuromeres, but during metamorphosis they develop extensive arborizations within the entire central nervous system. Nevertheless, there is no anatomical evidence that any such axons could release the peptide into the hemolymph.

Using transgenesis we expressed in the *Drosophila* SIFamide neurons either an apoptotic protein to kill them or RNAi to eliminate the peptide. Those experiments revealed that the four neurons and the peptide they produce are essential for the correct execution of male sexual behavior; in the absence of SIFamide males court males as intensely as females (Terhzaz et al., 2007). In *Drosophila* the fruitless gene produces male and female specific transcription factors that allow for the correct development of male and female brain archictecture and subsequent sexual behavior. The effects of SIFamide on male sexual behavior is at least in part through neurons expressing fruitless (Sellami and Veenstra, 2015).

More recent studies have started to look at other species, such as the blood feeding bug *Rhodnius prolixus*, the locust *Schistocerca gregaria* as well as several cockroaches (Ayub et al., 2020; Gellerer et al., 2015; Arendt et al., 2016). In all these species orthologs of the four neuroendocrine cells in the *pars intermedia* are the major SIFamide immunoreactive neurons. In cockroaches and *Schistocerca* there are other additional SIFamide immunoreactive neurons, but only in the cockroach *Rhyparobia maderae* and *Rhodnius* have SIFamide immunoreactive projections been found in the corpora cardiaca (Ayub et al., 2020; Arendt et al., 2016), suggesting an endocrine function for SIFamide in those species.

The deorphanization of the G-protein coupled receptor (GPCR) encoded by *Drosophila* gene CG10823 as the SIFamide receptor (Jørgensen et al., 2006) allowed the identification of SIFamide receptor orthologs from other species, such as one from the tick *Dermacentor variabilis* that is expressed in the male reproductive system (Sonenshine et al., 2011).

These initial data might suggest an important and perhaps preponderant role in reproduction and sexual behavior, but SIFamide and its receptor have now been been shown to be important for the correct execution of other behaviors as well, such as sleep in *Drosophila* and agression in a decapod crustacean (Park et al., 2014; Martelli et al., 2017; Dreyer et al., 2019; Vázquez-Acevedo et al. 2009), while in the tick *Ixodes scapularis* SIFamide innervates the salivary gland and in *Rhodnius* it is released during feeding and increases meal size as well as the rate of heart beat (Šimo et al., 2009; Ayub et al., 2020).

In some insect species the SIFamide gene has a paralog. IMFamide is a SIFamide paralog that was initially described from the silkworm *Bombyx mori* (Roller et al., 2008), but seems to be generally present in Lepidoptera. An independent SIFamide gene duplication occurred in the termite *Zootermopsis nevadensis* and the locust *Locusta migratoria* where the paralog peptide is SMYamide (Veenstra, 2014). Whereas the primary sequence of IMFamide shows significant differences from SIFamide, SIFamide and SMYamide peptides have remarkably similar structures. The recently published genome sequence for the American cockroach *Periplaneta americana* (Li et al., 2018) similarly reveals both a SIFamide and a SMYamide gene. I here report that in the latter species SMYamide is specifically expressed in two neurons innervating the salivary glands and that it seems likely that SMYamide also has a hormonal function.

## 2. Materials and Methods

### 2.1 Animals

*Periplaneta americana* are from a small colony that I maintain on mouse chow and water and that was intially established from animals obtained from Professor Peter Kloppenburg (University of Cologne, Germany). Only adults were used here.

### 2.2 RNA extraction and cDNA synthesis

Total RNA was isolated from specific tissues using a kit from Macherey-Nagel (Hoerdt, France). Moloney murine leukemia virus reverse transcriptase (New England Biolabs, Evry, France) and random hexamer primers were used to transcribe 1 μg RNA in a 20 μl reaction into cDNA.

### 2.3 RT-PCR for receptor expression

One microliter of cDNA was amplified by PCR using OneTaq DNA Polymerase (New England Biolabs) with with the following primers for the two putative SIFamide receptor: SIFaR1-forward 5’-CAGTCTCATTGCAGTCTCGC-3’, SIFaR1-reverse 5’-GCATCTTGACCACCTTGACC-3’, SIFaR2-forward 5’-CAACCTCTTCATCGCCAACC-3’, SIFaR2-reverse 5’-GCGGGAACAGATAGCACATG-3’, while actin, as a control, was amplified with primers actin-forward 5’-GCTATCCAGGCTGTGCTTTC-3’ and actin-reverse 5’-CAGGAAGGAAGGTTGGAACA-3’. All three primer pairs span an intron in the *Periplaneta* genome and yield products of around 400 bp. PCR profiles conisted of 30 sec denaturation at 94 °C, followed by 32 cycles of 15 sec at 94 °C, 15 sec at the annealing temperature and 30 sec at 68 °C. This was followed by final extension at 68 °C for three minutes. Annealing temperatures were 54 °C for the putative SIFamide receptors and 52 °C for actin. Bands were cut from the gel, purified and their identity confirmed by sequencing.

### 2.4 Immunohistology

Tissues were fixed in phosphate buffered 4% paraformaldehyde in Eppendorf tubes for 2 to 4 hrs at room temparature. After eight 30 minute washes in PBS containing 0.1% sodium azide and 1% Triton X100 (PBSAT), tissues were incubated for 1 hr in 10% normal goat serum (NGS) in PBSAT. Primary antisera diluted in 10% NGS in PBSAT were then added and tissues were incubated at room temperature for three days. Eight 30 minute washes with PBSAT and a 1 hour preincubation in 10% NGS in PBSAT preceded a two day incubation in secondary antiserum. This was followed by eight 30 minute washes in PBSAT after which tissues were transferred to small dissection dishes [to facilitate exchanging the glycerol solutions]. The procedure was continued with four 15 min changes in increasing concentrations of glycerol (20 %, 40 %, 60 % and 80 %) and finally tissues were mounted between a slide and coverslip in 80 % glycerol. All incubations at room temperature were performed on a gently rotating orbital shaker.

Two rabbit antisera were used, an old commercial antiserum to 5-hydroxytryptamine (5HT) from Immunotech (Marseille, France) and my own against SIFamide (Terhzaz et al., 2007). IgG from the SIFamide antiserum was isolated using capryllic acid and then coupled to tetramethyl-rhodamine to allow demonstration of both 5HT- and SIFamide-immunoreactivity in the same preparation. For such double labelings (Fig. 3), tissue was first incubated in the 5HT antiserum, followed by an fluorescein labeled Fab fragment of goat anti-rabbit IgG. The tissue was subsequently incubated in the rhodamin labeled SIFamide IgG for another two to three days. Primary antisera were diluted 1:4,000 (SIFamide) or 1:400 (5HT), the rhodamine labeled SIFamide antiserum was diluted 1:400. Secondary antisera were Alexa488-labeled goat anti-rabbit IgG and FITC-labeled Fab fragment from goat-anti rabbit IgG (Jackson Immunoresearch Europe Ltd, Ely, UK).

### 2.5 In situ *hybridization probes*

cDNA from brain-subesophageal complexes was used to amplify partial coding sequences for the SIFamide and SMYamide *Periplaneta* genes with Q5 polymerase (New England Biolabs), with the following primers: SIFa-forward 5’-TGTTGCCACATGTCTGCTTC-3’, SIFa-reverse 5’-GAAACCACGCTGAGCAGG-3’, SMYa-forward 5’-ATGAAATTCGCCTGCACCG-3’ and SMYa-reverse 5’-GCAGACCTCTACAGCCATCT-3’. PCR products were gel-purified and quantitated and then used to make digoxigenin-labeled anti-sense probes using single primer PCR with Taq polymerase (New England Biolabs) in which a third of the dTTP had been replaced with Digoxigenin-X-(5-aminoallyl)-2’-deoxyuridine-5’-triphosphate (Jena Bioscience, Jena, Germany) for 40 cycles. The final PCR product was used without purification and diluted ten times with HS for *in situ* hybridization experiments.

### 2.6 Combined in situ *hybridization and immunohistology*

The *in situ* hybridization protocol is based on a protocol for *Drosophila* (Kim et al., 2006) and was modified in order to take into account the much larger size of the cockroach CNS. Dissections were done in 0.9 % NaCl and tissues were fixed in phosphate buffered 4% paraformaldehyde for 2 to 4 hrs at room temperture in Eppendorf tubes. Tissues were subsequently washed thrice for 30 min in PBS with 0.2 % Tween 20 (PBST) and once for 30 min in 70% ethanol. The 70% ethanol was then changed and the tissues stored at -20 °C usually for three days, but sometimes longer. Tissues were next washed thrice for 30 min in PBST and then incubated with Proteinase K (25 µg/ml) in PBST for 45 min. The proteinase K was stopped by washing the tissue for 30 min in glycin (2 mg/ml) in PBST and this was followed by two washes of 30 min in PBST. The tissues were then fixed a second time in phosphate buffered 4% paraformaldehyde for 1 hr followed by two washes in PBST for 30 min and one for 30 min in 1:1 mixture of hybridization solution (HS: 50 µg/ml heparin, 100 µg/ml salmon testes DNA, 750 mM NaCl, 75 mM sodium citrate, in deionized RNase free water, pH 7.0, 50% deionized formamide, 0.1 % Tween 20) and PBST. Next the solution was replaced with HS and incubated for 30 min in HS. Up to this point everything is done at room temperature. HS is then replaced with fresh HS and the tissues are put in a water bath at 48 °C. Two hours later HS is replaced with HS containing up to 10% of digoxygenin hybridization probe which has previously been brought to 95 °C for either 3 min, in case of a previously used probe, or 45 min when a hybridization probe is used for the first time. Hybridization is carried out overnight at 48 °C. The following day hybridization probe is recovered and stored at -20 °C for future use. Tissues are washed in fresh HS five times, first three times for 2.5 to 4 hrs, then once overnight, and a final wash for 3 to 4 hrs; all washes are at 48 °C. The tissues are then removed from the water bath and the remainder of the procedure is performed at room temperature. The HS is replaced with the 1:1 mixture of hybridization solution for 30 minutes, followed by three washes in PBST for 30 min each after which the tissues are saturated with 1% BSA in PBST for 1 hr. Next the 1% BSA in PBST is replaced with the same solution to which both sheep anti-digoxygenin Fab-fragments conjugated to alkaline phosphatase (Roche Diagnostics GmBH, Mannheim, Germany) in a 1:1,000 dilution and SIFamide antiserum in a 1:2,000 dilution have been added. Incubation is overnight at room temperature. Next morning tissues are washed twice 30 min with PBST, followed by three washes in freshly prepared alkaline phosphate buffer (APB, 100 mM Tris, 50 mM MgCl2, 100 mM NaCl, pH 9.5 and 0.1% Tween 20). During the last wash, the tissues are transferred to small glass dissection dishes and after the last wash, the digoxygenin probe is visualized by replacing the APB is replaced with the same containing 20 µl/ml of a commercial NBT/BCIP stock solution (Roche Diagnostics GmBH, Mannheim, Germany, 18.75 mg/ml nitro blue tetrazolium chloride and 9.4 mg/ml 5-bromo-4-chloro-3-indolyl-phosphate, toluidine-salt in 67% DMSO). Color development is followed under a binocular and may take 5 to 15 minutes and once it is judged satisfactory, the alkaline phosphatase is stopped by changing the staining solution with 100 mM phosophate buffer, pH 7.0. Note that from now on no detergent can be used, as it may dissolve the product from the alkaline phosphatase reaction. After two washes of 5 minutes with the 100 mM phosphate buffer, tissues are transferred to an Eppendorf tube, followed by five 30 min washes in PBS containing 0.1% sodium azide (PBSA). Tissues are then incubated in 10% normal goat serum in PBSA, followed by incubation over night with secondary antisrum diluted 1:1000 in 10 % normal goat serum in PBSA at room temperature. The next day, the tissues are washed eight times for 30 min in PBSA and then transferred to small dissection dishes in which they are exposed to increasing concentrations of glycerol (20 %, 40 %, 60 % and 80 %, 15 minutes each) and then tissues are mounted in 80 % glycerol.

### 2.7 Bioinformatics

Methods employed in bioinformatics have been described in detail in a previous manuscript (Veenstra, 2020). For the expression of SIFamide, SMYamide and their putative receptors in *Periplaneta* the followoing transcriptome short read archives (SRAs) were analyzed: DRR014884, DRR014885, DRR014886, DRR014887, DRR014888, DRR014889, SRR921630, SRR5286150, SRR5286151, SRR5286152, SRR5286153, SRR5286154, SRR1184457, SRR1184458, SRR1322009, SRR2994649, SRR2994650, SRR3056857, SRR3056858, SRR3089536, SRR3089537, SRR3089538, SRR3289663, SRR3289684, SRR3289687, SRR5097509, SRR5097510, SRR5097511, SRR5097512, SRR5097513, SRR5097514, SRR5097515 and SRR5097516. These were downloaded from NCBI: https://www.ncbi.nlm.nih.gov/sra.

Coding sequences for SIFamide and SMYamide precursors as well as those for putative SIFamide receptors were deduced from transcriptome and genome data as described (Veenstra, 2020). All these sequences are listed in the supplementary spreadsheet. Sequence logos for the neuropeptides were made from these peptide using https://weblogo.berkeley.edu/. A phylogenetic tree from the SIFamide GPCRs was made by using clustal omega (Sievers et al., 2011) to align the sequences and Fasttree2 (Price et al., 2010) to produce a tree from of the alignment of their transmembrane regions.

## 3. Results

### 3.1 SMYamide distribution and sequences

SMYamide precursors were found in several Polyneoptera insect orders. The SIFamide gene duplication seems to have occurred after the Plecoptera diverged, since SMYamide genes have been found in Orthoptera, Embioptera, Phasmatodea, Mantodea and Blattodea, but could not be found in genomes from Plecoptera or Polyneopteran insect orders that evolved even earlier. Analysis of Polyneopteran transcriptome yielded similar results (Bläser and Predel, 2020). The structure of the various SMYamide peptides is not as well conserved as that from the SIFamides (Figs. 1, S1, S2).

**Fig. 1.**
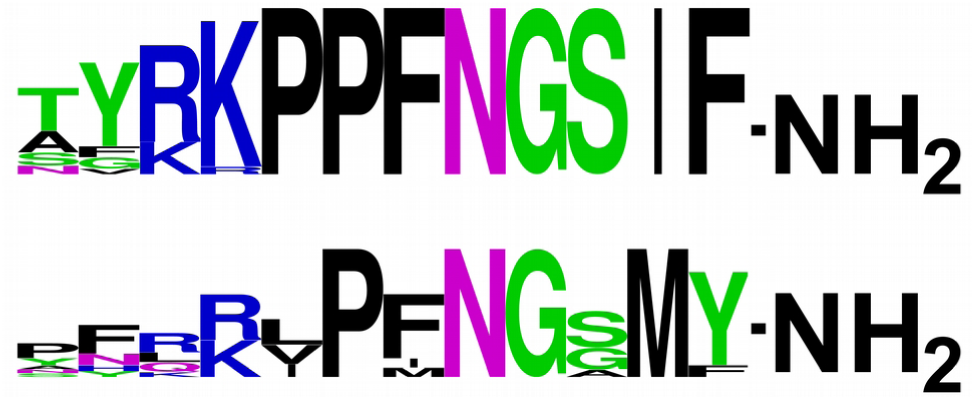
Sequence logos of SIFamide and SMYamide. Note that the SIFamide peptide sequences are better conserved than the SMYamide sequences.

### 3.2 SIFamide-immunoreactivity in Periplaneta

SIFamide immunoreacitivity in the brain of *Periplaneta americana* has been previously described in detail and six different SIFamide immunoreactive cell groups were identified (Arendt et al., 2016). Although I used whole mounts rather than sections which may make it more difficult to identify neurons, the same cells were found here, although group 6 was not always identified. These cells have all been previously described and so there is no need to that here. In the ventral nerve cord all ganglia show extensive SIFamide immunoreactive arborizations of the four large interneurons in the brain as also described for the locust *Schistocerca gregaria* (Gellerer et al., 2015). When SIFamide immunoreactive neurons were encountered in thoracic or abdominal ganglia there were not symmetrical, their axons could only be followed for a short distance into the neuropile and they seemed to be local interneurons (Fig. 2c). As in *Schistocerca* (Gellerer et al., 2015) no SIFamide immunoreactive material was detected in any of the efferent nerves from these ganglia.

**Fig. 2.**
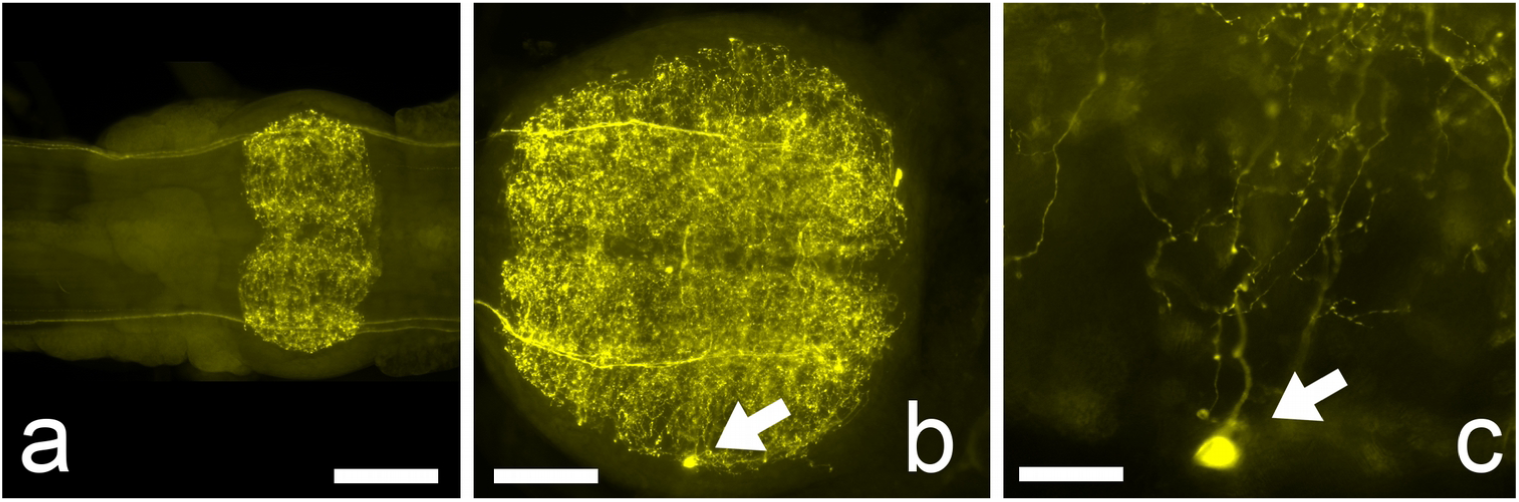
SIFamide immunoreactivity in the fourth abdominal (a) and terminal abdominal ganglia (b) of a male cockroach. On occasion one can find small SIFamide immunoreactive neurons like the one in the terminal ganglion (arrow in b) at a higher magnification it is possible to follow their axons for a short distance (arrow in c). Scale bars 200 µm (a and b) and 50 µm (c).

However, two prominent SIFamide immunoreactive neurons were found in the subesophageal ganglion (Fig. 3). Their axons leave the ganglion through the salivary duct nerve (SDN) that is known to contain two large axons from the contralateral salivary neuron 1 (SN1), the ipsilateral salivary neuron 2 (SN2) as well as a few smaller 5HT-immunoreactive neurons. SIFamide and 5HT immunoreactivities do not colocalize (Figs 3b,c). Both the position within the ganglion and the ipsilateral projections confirm that the SIFamide immunoreactive neurons are the SN2’s. The innervation of salivary glands by the SIFamide immunoreactive axons appears superficial, as the axons seem to surround the salivary acini, rather than making close contact with them like the 5HT axons that terminate between the lateral membranes of inidividual cells of the acini (Fig. 3b) and that also reach the gland through the SDN.

**Fig. 3.**
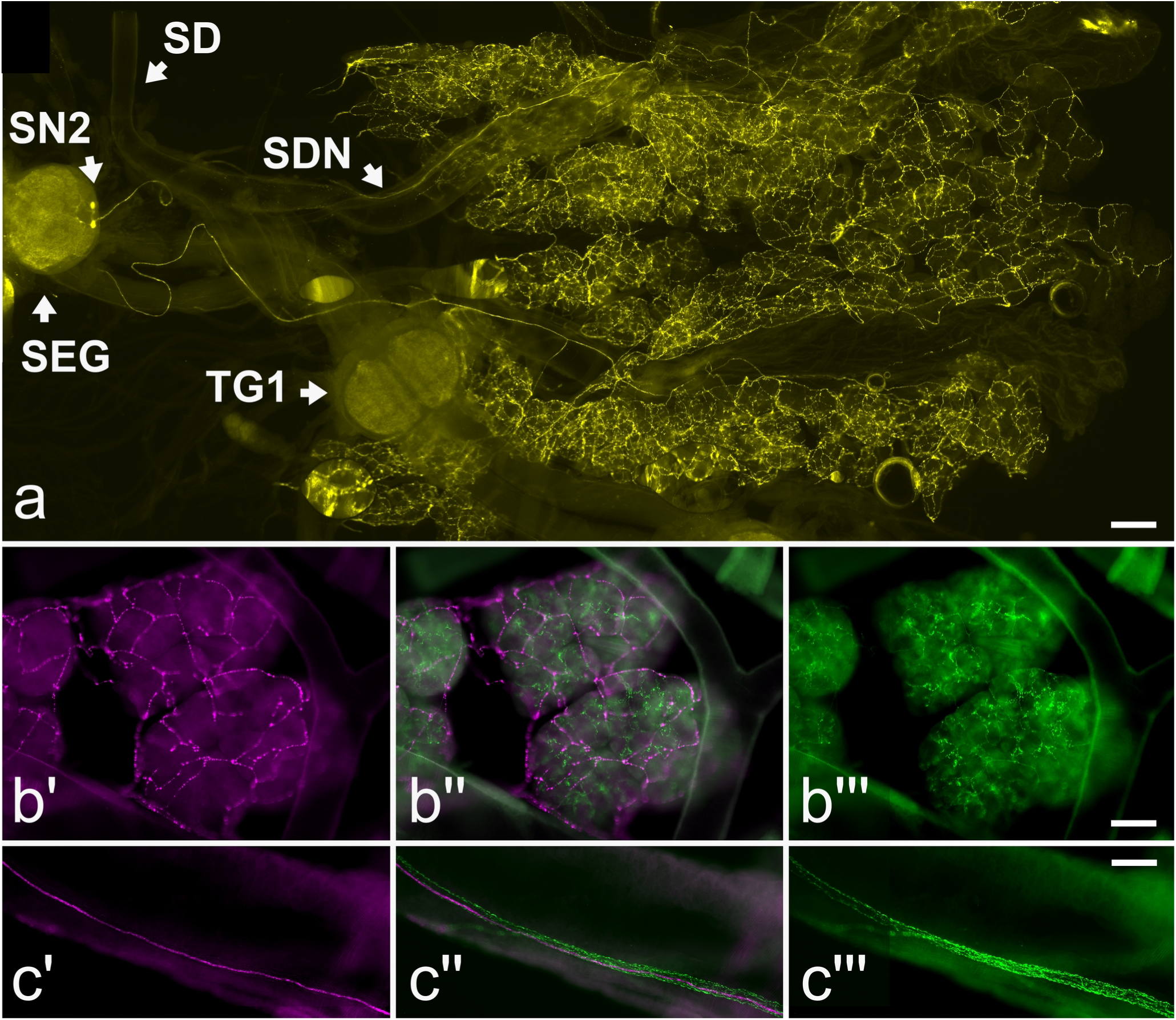
SIFamide immunoreactive neurons innervating the salivary gland. In (a) an overview of part of the ventral nerve cord with the subesophageal ganglion (SEG) and the first thoracic ganglion (TG1), the salivary gland (the structure that occupies the right half of the picture) and the salivary duct (SD) as well as the salivary duct nerve (SDN) through which the axons of the two salivary neurons 2 (SN2) run to the salivary gland. Note the extensive branching of the two SN2 neurons inside the salivary gland. In (b) SIFamide immunoreactive innervation in individual acini is revealed by magenta (b’ and b’’), while 5HT-immunoreactive axons are in green (b’’ and b’’’). Note that the innervation by SIFamide appears very superficial, while the 5HT-immunoreactive axons penetrate between the individual cells of the acini. In (c) the single SIFamide axon in the SDN shown in magenta (c’ and c’’) is accompanied by a number of smaller 5HT-immunoreactive axons in green (c’’ and c’’’). Scale bars 400 µm (a) and 100 µm (b and c).

As previously suggested by others (Baumann et al., 2004) these observations suggest that the axon terminations of the SN2’s within the salivary glands represent neurohemal release sites. It can thus be expected that significant quantities of peptide will be released into the hemolymph when these neurons are active and that the peptide acts as a hormone on other target tissues as well as on the salivary gland. Thus one question of interest concerns whether receptors might be expressed in other tissues. As the cockroach has two SIFamide-like genes, there is also the question which of the two peptides is produced by the SN2 neurons.

### 3.3 In situ *hybridization*

Coding sequences for SIFamide and SMYamide are very short and this limits the choice of suitable primers for PCR amplification. Consequently, the hybridization probes are also short and the signals are not very strong. Nevertheless, *in situ* hybridization yielded positive signals for the SIFamide antisense probe in the large SIFamide interneurons in the brain as well as in some neurons of cell groups 2 and 3. In the subesophageal ganglion no reliable SIFamide hybridization signals were found, but the two SN2 neurons yielded a positive signal for SMYamide (Fig. 4).

**Fig. 4.**
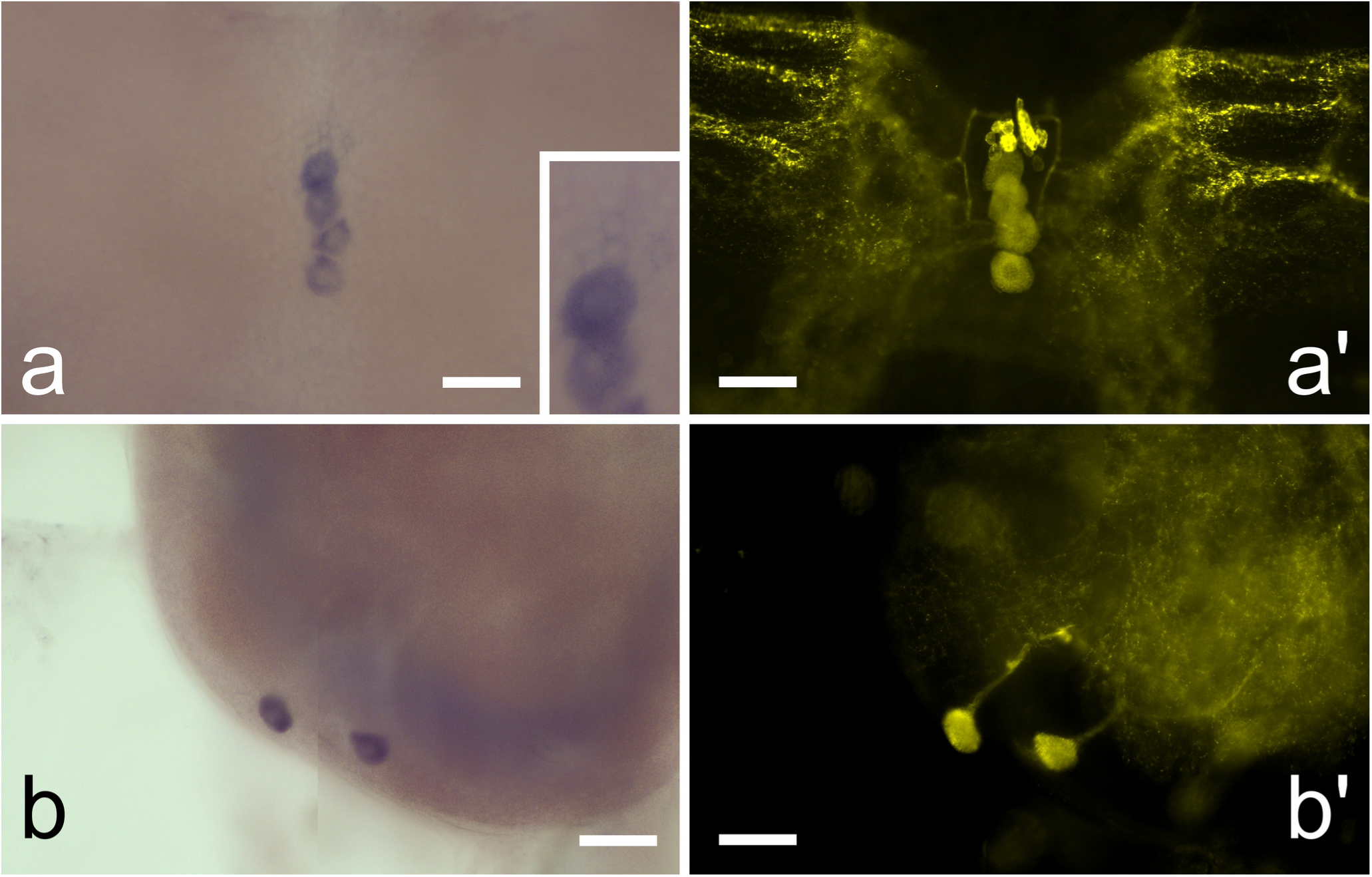
*In situ* hybridization. In (a) SIFamide *in situ* hybridization localizes to the four big SIFamide immonreactive interneurons of the brain identified by immunofluorescence in (a’). The much smaller SIFamide immunroeactive neurons on top of the larger ones are also recognized by the SIFamide anti-sense probe, although very weakly (insert in a’). In (b) SMYamide *in situ* hybridization signal (b) localizes to the SN2 neurons of the subesophageal ganglion as shown by simultaneous labeling of these neurons with SIFamide antiserum (b’). All scale bars 100 µm.

These results are consistent with transcriptome analyses from brain and subesophageal ganglia in this species where 13,852 SIFamide and 34 SMYamide specific reads were found in brain SRAs versus 1,122 SIFamide and 44,733 SMYamide specific reads in SRAs from the subesophageal ganglion (for details see Table S1).

### 3.4 SIFamide receptors

As in the termite *Zootermopsis* (Veenstra, 2014) the *Periplaneta* genome contains two genes coding for a SIFamide-like receptor. There are three possible explanation for the existence of two such receptors in these species. First, these species have both a SIFamide and a SMYamide gene and it is plausible that the two neuropeptides each have their own specific receptor. Secondly, the second putative SIFamide receptor could have another as yet unknown ligand and lastly, both receptors might be activated by SIFamide and SMYamide.

A comparative sequence analysis of arthropod SIFamide receptors yields a phylogenetic tree that reveals two major branches (Fig. 5). One branch contains the deorphanized *Drosophila* and *Bombus terrestris* SIFamide receptors (Jørgensen et al, 2006; Lismont et al., 2018), the other the deorphanized *Ixodes*-2 SIFamide receptor (Šimo et al., 2013). Furthermore, in the planthopper *Nilaparvata lugens* no SMYamide gene can be found, but it does have two SIFamide receptors (Tanaka et al., 2014). It thus appears likely that all these GPCRs are indeed activated by both SIFamide and SMYamide. The position of the deorphanized *Drosophila* SIFamide receptor on the phylogenetic tree is unusual and a close inspection of the various arthropod SIFamide receptor sequences shows the sequence from *Drosophila* to differ significantly from the others (Fig. S3).

**Fig. 5.**
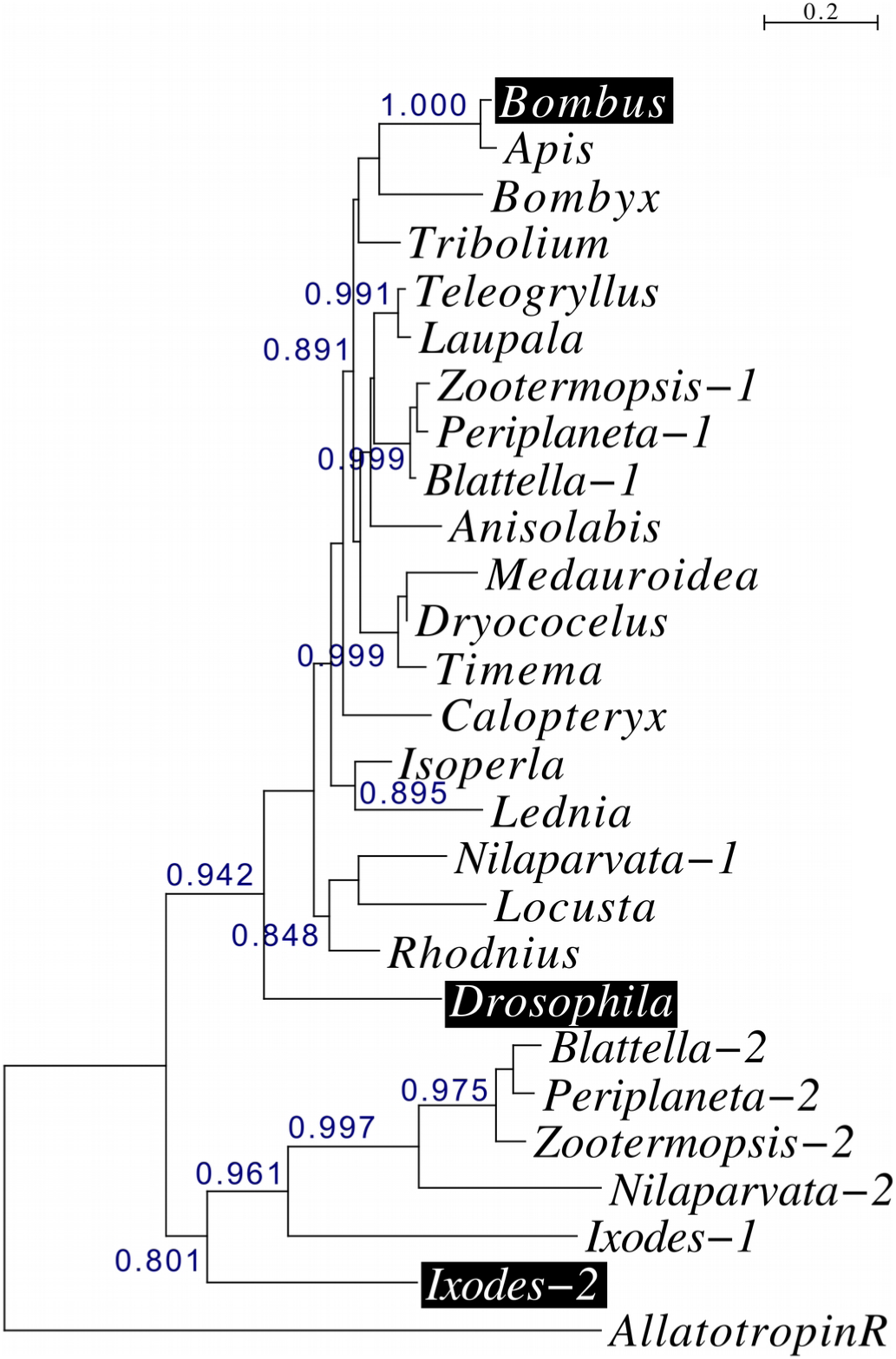
Phylogenetic tree of deorphanized arthropod SIFamide receptors, highlighted in black, and their orthologs from a number of additional species. Note that the tree reveals two major branches, each of which contains at least one deorphanized receptor. Branch probabilities of at least 0.80 have been indicated on the tree that was rooted on the allatotropin receptor from *Schistocerca gregaria*.

### 3.5 SIFamide receptor expression in Periplaneta

PCR on cDNA from a variety of tissues show that both putative SIFamide receptors are expressed not only in the central nervous system and the salivary glands, but also in the testis and ovary and a low level expression seems to be present in the midgut, Malpighian tubules, fat body and flight muscle (Fig. 6). Sequence analysis of the purified PCR products confirmed unambiguously the identities of the PCR products (Figs. S4 and S5).

**Fig. 6.**
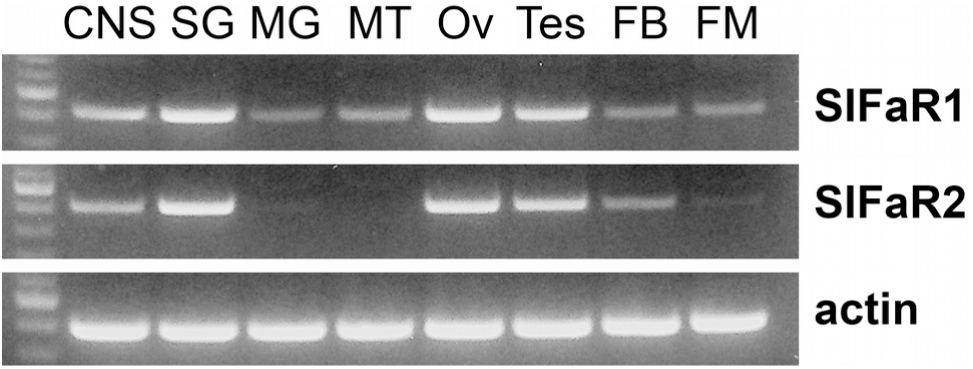
RT-PCR of the two *Periplaneta* SIFamide receptor-related GPCRs and actin in various tissues. First lane in each gel contains a 100 bp DNA ladder; the more intense band corresponds to 500 bp. CNS, central nervous system; SG, salivary gland; MG, midgut; MT, Malipighian tubules; Ov, ovary; Tes, testes; FB, fatbody; FM, flight muscle. Note the significant expression in the CNS, salivary glands and the gonads and minor expression in other tissues.

The number of transcriptome SRAs from Periplaneta is relatively small and covers only some of the tissues. Nevertheless it is interesting to note that testes SRAs confirm the presence of SIFamide receptor 1 specific reads while the abundance of SIFamide receptor reads in whole body extracts from females can not explained by their expression in the central nervous system and thus indirectly suggests expression of these receptors in peripheral tissues (Table S1).

## 4. Discussion

This study reveals that two previously identified neurons in the subesophageal ganglion that innervate the salivary gland are immunoreactive with antiserum to SIFamide. *In situ* hybridization experiment show that these neurons express the SMYamide rather than the SIFamide gene. Publicly available expression data from the brain and subesophageal ganglion support this conclusion. Results also show that the SIFamide receptor is expressed in tissues other than the brain and the salivary glands.

It is not surprising that the SIFamide antiserum recognizes SMYamide, as the two peptides show significant sequence similarity with C-termini that are almost identical (Fig. 1). Isoleucine-methionine and phenylalanine-tyrosine are both common amino substutions in neuropeptide evolution and the remainder of the sequences are also similar. In *Nauphoeta cinerea*, a different cockroach species, the SN2 neurons were shown to contain the large electron dense granules that typically contain neuropeptides (Maxwell, 1978). Indeed it had been previously speculated that these neurons also produce a neuropeptide in *Periplaneta americana* (Baumann et al, 2004). An earlier paper showed the SN2 neurons be immunoreactive with GABA antisera and this is thus a likely case of co-localisation of a classical neurotransmitter with a neuropeptide (Rotte et al., 2009).

The SN2 neurons in the cockroach have been well characterized. Electrophysiological recordings from this nerve during feeding show two large action potentials, with the largest one derived from the dopaminergic salivary neuron 1 and the smaller of the two from SN2, as well as a number of much smaller action potentials believed to derive from the 5HT-immunoreactive axons (Watanabe and Mizunami, 2006). In the cockroach the cell bodies from which these 5HT axons originate are unknown, but similar neurons in locusts have been called satelite neurons and are located in the subesophageal ganglion (Schachtner and Bräunig, 1995) and is seems plausible that these are homologs of the SEN neurons in *Periplaneta* (Davis, 1987).

Dopamine is known to stimulate fluid secretion but does not seem to stimulate release of protein, while 5HT stimulates secretion of both fluid and a variety of proteins (Just and Walz, 1996), which one assumes represent digestive enzymes. The effects of SMYamide on the salivary gland remain to be determined. GABA, which is co-localized with SMYamide in SN2, alone has no effect on either fluid secretion or electrical activity of the salivary gland neurons, but it reinforces both these parameters when the salivary duct nerve is electrically stimulated (Rotte et al., 2009). Since such stimulation must also lead to release of SMYamide one has to assume that SMYamide stimulates salivation, even though it is unknown whether this concerns the secretion of fluid, digestive enzymes, or both.

The SMYamide gene has its origin in a duplication of the SIFamide gene. There are different reasons as to why duplicated neuropeptide may not be not selected against and thus survive during evolution. The need for a high quantities of gene products is one of them. This is likely the case for the amplified vasopressin-like genes in *Locusta migratoria*, where an initially peptidergic interneuron evolved into a neuroendocrine cell and therefore almost certainly needs to be able to produce much larger quantities of peptides. One might be tempted at first sight to think that the very large SIFamide interneurons in the brain might similarly benefit from a second gene producing a very similar peptide. However the absence of significant expression of SMYamide in the brain suggests that this is not the case. Another scenario is the specialization of the different genes for different functions, the expression of SIFamide in the brain interneurons and SMYamide by the neurons in the subesophageal ganglion shows that this could be the case here.

Based on the pattern of innervation of salivary glands by the SN2 neurons it has previously been suggested that these neurons are neuroendocrine in nature (Baumann et al., 2004). The expression patterns of the two *Periplaneta* SIFamide receptors reinforces the notion that SMYamide is indeed a hormone. The first SIFamide was identified using contractions of the locust oviduct as a bioassay (Janssen et al., 1996), but work on flies failed to find any evidence for the possible release of this peptide into the hemolymph. It was neither possible to find SIFamide innervation of the oviduct or any other potential target tissue. Although the *Drosophila* SIFamide receptor is expressed outside the nervous system, it is in afferent neurons that have axons projecting into the nervous system where they are exposed to SIFamide (Sellami and Veenstra, 2015). The data presented here show significant expression of both SIFamide receptors in peripheral tissues, particularly the gonads. In the bumblee the SIFamide receptor is also expressed in the testes as well as other peripheral tissues (Elsmont et al., 2018). This suggests that perhaps these peptides, *i.e*. SIFamide, SMYamide and/or IMFamide, may function as endocrines in other species as well, including holometabolous insect species. The phylogenetic tree of the SIFamide receptor is intriguing in this respect. The *Drosophila* receptor looks like the most ancient one in the first branch of this tree (Fig. 5). Obviously, this is not the case, as it is one of the most recent ones to evolve. This suggests that there is less evolutionary pressure on maintaining its primary sequence in the fruit fly than there is in the other insect species. The affinity of neuropeptide-GPCR interactions are typically much smaller when the neuropeptide is only released inside the nervous system. From experiments where SIFamide or the neurons it produces were eliminated, it is clear that the peptide continues to play an important role in this species (Terhzaz et al., 2007), but in *Drosophila*, and likely all flies, SIFamide has no endocrine role and only retains a function within the central nervous system. The loss of an endocrine function could thus be the reason why there is less pressure on maintaining the structure of its receptor. Hence the phylogenetic tree of the SIFamide receptors also points to the possibility that these peptides could have endocrine functions in holometabolous insects.

The effects of SIFamide on the salivary gland have not yet been studied, but as the SN2s are active during feeding one may assume that it stimulates the production of saliva. In adult cockroaches increased feeding will generally lead to increased production of egss and sperm. So the activation of the SIFamide receptors in the gonads and the fat body while the insect feeds may prime the reproductive organs as well as the fat body for the nutrients that are about to arrive.

It are the pioneer neurons that produce SIFamide (Gellerer et al., 2015). It is perhaps not unreasonable to expect that the first neurons that are needed in a developing embryo are those that make sure the animal will be able to feed. The coincidence that the same peptide might stimulate salivation in both blood feeding ticks and cockroaches and is also expressed in pioneer neurons, is therefore intriguing. Natural selection favors survival of species, for a species to survive its individual members need to survive first. Once individuals are well fed and able to produce viable gametes they should reproduce. If SIFamide and SMYamide were peptides that regulate feeding in a broad sense, as seems plausible (Martelli et al., 2017; Dreyer et al., 2019), it is perhaps not surprising that once they are no longer released (or eliminated by genetic manipulation) insects increase their sexual behavior. This might help explain why elimination of SIFamide in *Drosophila* leads to hypersexual behavior.

