## SupplementaryData for "The neuropeptide SMYamide, a SIFamide paralog, is expressed by salivary gland innervating neurons in the American cockroach and likely functions as a hormone"

Jan A. Veenstra

### Content

Fig. S1. Deduced amino acid sequences of various Arthropod SIFamide precursors. 2

Fig. S2. Deduced amino acid sequences of various Polyneopteran SMYamide precursors. 3

Table S1. Analysis of SIFamide-related gene expression in *Periplaneta* transcriptome SRAs. 4

Fig. 3. Sequence alignment of various SIFamide GPCR homologs. 5

Fig. S4. Sanger sequence analyis of the RT-PCR product for *Periplaneta* SIFamide receptor 1. 12

Fig. S5. Sanger sequence analyis of the RT-PCR product for *Periplaneta* SIFamide receptor 2. 13

***Anisolabis maritima***
MQKSSSLLLLIVAVALLLILEDGALAYRKPPFNGSIFGKRGSSAADFDAAVKALSSMCEIATEACSAWFPQNESNN*

***Apis mellifera***
MVSTRLVLAVVAALFVLAISVDAAYRKPPFNGSIFGKRSNTITDYEITSRAMSSVCEVVSETCNAWLSRQDSN*

***Blattella germanica***
MQNRVAATCLLLLAVLLFADLAAATYRKPPFNGSIFGKRGNVVEYDGTGKALSALCEIASEACSAWFPSADNN*

***Bombus terrestris***
MSSRFVVAIVVALFILAIAVDAAYRKPPFNGSIFGKRSNAITDYELTSRAMSSVCETVSETCNAWLARQDSN*

***Bomyx mori***
MRADLIYFMFLVIILTLATIEATYRKPPFNGSIFGKRNNVENDSSGRAIAALCEITTETCQAWYQALESQ*

***Calopteryx splendens***
MNSKAIFIFAVVLVAFALLSGSADASSYRKPPFNGSIFGKRGSGASADAESRGVAFSALCEVAIEACSSWMPASQDSK*

***Drosophila melanogaster***
MALRFTLTLLLVTILVAAILLGSSEAAYRKPPFNGSIFGKRNSLDYDSAKMSAVCEVAMEACPMWFPQNDSK*

***Dryococelus australis***
MQRSGVLICAVMLAVLLLLSEPAAATGKKPPFNGSIFGKRSNYVAEIRDYESTGRSLSAMCEIASEACASWFPQADKR*

***Isoperla grammatica***
MQKSGVATCLLVLVLLLASDFAAANYRKPPFNGSIFGKRVGNSADYDGAAKALSSMCEIAQEACAAWFPQAEASK*

***Ixodes scapularis***
MNSWKAFFMFGTLLVMAVMMNMACAAYRKPPFNGSIFGKRSRADLNNADVKYAMCEAVWDTCTQWFPITQDGAQ*

***Laupala kohalensis*** (incomplete)
LRFAAVLCVALLLVAASAPSAASYRKPPFNGSIFGKRASAAGAAAAPTNAQEYEAAAAGKALSALCEVAAEACSAWFPQQNELN*

***Lednia tumana***
MQKSALVKCIFLLALLAVCAEITTATYRKPPFNGSIFGKRAGISIDYDNAGKALSSMCEIATEACAVWFPQRDESK*

***Locusta migratoria***
MQTAACSRFLVVVVLALVMFTAAASAAAATFRRPPFNGSIFGKRNSIESAGSSTAVAAVCEIAAEACAAWLNNDK*

***Medauroidea extradentata***
MQRSGVIVCVVLVALLLLLSEPAVATGKKPPFNGSIFGKRSNYVAELRGCRGVVIADYESTGRSLSAMCEIASEACASWFPQADKR*

***Nilaparvata lugens***
MVRICLAMCILVSIILAFVDAAYKKPPFNGSIFGKRGSTIIEYENAGKALSSLCEIASEACSAWFPLPDN*

***Periplaneta americana***
MQTRGVATCLLLLAVLLLAEFAAATYRKPPFNGSIFGKRGNAADYDGAGKALSAMCEIASEACSAWFPQADNN*

***Rhodnius prolixus***
MSRTLFVCCFTLVVALIFLDAAMATYKKPPFNGSIFGKRAGPSSDYETAGKALSTMCEIAAEACSAWFPVQDNN*

***Teleogryllus comodus***
MAGKSFAASRRSARPLAVLLRAAAVLCVALLLVAASAPSAASYRKPPFNGSIFGKRAPAAAAPAPAASAQDYEAAAAGKALSALCEVA
AEACSAWFPQQNELN*

***Timema cristinae***
MNKYSIARCMLLLTLFMLTELTLANVKKPPFNGSIFGKRGNYGGELRDYDSAGRTLSVMCEIASEACSGWFPQPDKR*

***Tribolium castaneum***
MQLALAKVFSVCIVVIILTSWIEMTEATYRKPPFNGSIFGKRGATIEYDSASKALSAMCEIASEACQTWFPSQEK*

***Zootermopsis nevadensis***
MQNRVVATCVLLLAVLLLAEFATAAFRKPPFNGSIFGKRGSPTDYDGASKALSAMCEIASEACSAWFPQMD*

**Fig. S1.** Deduced amino acid sequences of various Arthropod SIFamide precursors. Coding sequences are present in the supplementary spreadsheet of this paper.

***Blattella germanica***
MQFSQSVIFFLAILLLTLSTTCNPGVPFRRLPFNGSMYGKRASSALPMDYDNNKAFSSLCELAAEVCSTWYPQQVENN*

***Dryococelus australis***
MLHKFALAIMIFACVLCVALAYNLRVPINGGMYGKRDGPSEYDARSKAVSTMCEMATEVCSAWLAQADPN*

***Laupala kohalensis*** (incomplete)
-------LAIGFALHPVLASGPFKKIPFNGSMYGKRASPSEYEAAAAGKALSALCEVAAEACSAWFPQQNELN*

***Locusta migratoria***
MNRCSLVAMLVWAILLQTCLTEGIAFQKLPFNGAMYGKRTTSVDFDSSNRAISSLCETASEVCASWYGQPDTN*

***Medauroidea extradentata***
MIHISIARSLLVLVCTLLVTLGYNLRVPMNGGMYGKRDGPSEYDARGKAVSTMCEMATEVCSAWLAQADPN*

***Periplaneta americana***
MKFACTVLSLVALLLLAVLVACNPGPPFRRLPFNGSMYGKRTGNALPMDYDSNSKALSSLCEMAVEVCPTWFPQQQENN*

***Teleogryllus comodus***
MTEASTPAATAWCVRLLCLLLLVAALLEAAAARGSFQKIPFNGSMFGKRAAATEYELSHALSAMCEAAAAACTAWTQPDK*

***Timema cristinae***
MILTAVLIALSQNAMAANYRKLPFNGGMYGKRDTNLGELLGPQILTVRAKLCHPCVRSPLKHAQCGFHSRRSTRATSDDLPR*

***Zootermopsis nevadensis***
MKLSCSMIFLLALLLALLVDCNTGPPHRRVPFNGSMYGKRTANSLSVDYDSNAKSLSSLCEVATEVCSAWFPQQTENN*

**Fig. S2.** Deduced amino acid sequences of various Polyneopteran SMYamide precursors. Coding sequences are present in the supplementary spreadsheet of this paper.

| SRA | Spots | SIFamide | | SMYamide | | SIFaR1 | | SIFaR2 | | |
|---|---|---|---|---|---|---|---|---|---|---|
| DRR014884 | 9576533 | 47 | **4.9** | 4 | **.4** | 0 | **.0** | 2 | **.2** | Egg cases |
| DRR014885 | 9442758 | 27 | **2.9** | 1 | **.1** | 0 | **.0** | 4 | **.4** | young larvae |
| DRR014886 | 9060819 | 5 | **.6** | 2 | **.2** | 4 | **.4** | 7 | **.8** | An old-instar female larva |
| DRR014887 | 9962367 | 1 | **.1** | 4 | **.4** | 2 | **.2** | 2 | **.2** | An old-instar male larva |
| DRR014888 | 8067940 | 5 | **.6** | 0 | **.0** | 0 | **.0** | 2 | **.2** | Adult female |
| DRR014889 | 9773451 | 5 | **.5** | 0 | **.0** | 0 | **.0** | 0 | **.0** | Adult male |
| SRR921630 | 10494946 | 0 | **.0** | 1 | **.1** | 2 | **.2** | 0 | **.0** | Generic sample |
| SRR1184457 | 27531950 | 0 | **.0** | 0 | **.0** | 0 | **.0** | 0 | **.0** | no specific information provided |
| SRR1184458 | 27531950 | 0 | **.0** | 0 | **.0** | 0 | **.0** | 0 | **.0** | no specific information provided |
| SRR1322009 | 38573631 | 3 | **.1** | 0 | **.0** | 64 | **1.7** | 0 | **.0** | testis |
| SRR2994649 | 26044067 | 3 | **.1** | 4 | **.2** | 0 | **.0** | 7 | **.3** | control whole body |
| SRR2994650 | 26565524 | 16 | **.6** | 6 | **.2** | 5 | **.2** | 41 | **1.5** | E.coli treated whole body pooled male and female |
| SRR3056857 | 25780592 | 3 | **.1** | 4 | **.2** | 0 | **.0** | 7 | **.3** | Whole body pooled male and female |
| SRR3056858 | 26236239 | 16 | **.6** | 6 | **.2** | 5 | **.2** | 40 | **1.5** | E.coli treated whole body pooled male and female |
| SRR3089536 | 22954896 | 1 | **.0** | 0 | **.0** | 1 | **.0** | 0 | **.0** | chemosensory organs |
| SRR3089537 | 21727373 | 4 | **.2** | 0 | **.0** | 0 | **.0** | 2 | **.1** | chemosensory organs |
| SRR3089538 | 21585983 | 52 | **2.4** | 107 | **5.0** | 2 | **.1** | 0 | **.0** | chemosensory organs |
| SRR3289663 | 27096583 | 0 | **.0** | 0 | **.0** | 4 | **.1** | 12 | **.4** | Foregut |
| SRR3289684 | 33591931 | 0 | **.0** | 0 | **.0** | 10 | **.3** | 0 | **.0** | Midgut |
| SRR3289687 | 32043575 | 0 | **.0** | 0 | **.0** | 2 | **.1** | 7 | **.2** | Foregut withTreatment by Cycloxaprid |
| SRR5097509 | 33490493 | 4088 | **122.1** | 2 | **.1** | 16 | **.5** | 28 | **.8** | Brain stung |
| SRR5097510 | 100881736 | 7369 | **73.0** | 32 | **.3** | 42 | **.4** | 2 | **.0** | Brain |
| SRR5097511 | 23403587 | 701 | **30.0** | 0 | **.0** | 12 | **.5** | 0 | **.0** | Brain |
| SRR5097512 | 41305288 | 96 | **2.3** | 4569 | **110.6** | 4 | **.1** | 0 | **.0** | SEG |
| SRR5097513 | 46825733 | 212 | **4.5** | 6279 | **134.1** | 0 | **.0** | 0 | **.0** | SEG |
| SRR5097514 | 132099525 | 556 | **4.2** | 22945 | **173.7** | 0 | **.0** | 0 | **.0** | SEG stung |
| SRR5097515 | 40310873 | 258 | **6.4** | 10940 | **271.4** | 0 | **.0** | 0 | **.0** | SEG stung |
| SRR5097516 | 24170647 | 1694 | **70.1** | 0 | **.0** | 0 | **.0** | 10 | **.4** | Brain stung |
| SRR5286150 | 43339233 | 12 | **.3** | 10 | **.2** | 2 | **.0** | 2 | **.0** | male adult whole body |
| SRR5286151 | 34455335 | 61 | **1.8** | 7 | **.2** | 14 | **.4** | 28 | **.8** | female adult whole body |
| SRR5286152 | 38897702 | 17 | **.4** | 0 | **.0** | 7 | **.2** | 14 | **.4** | newly molted whole body |
| SRR5286153 | 64343011 | 51 | **.8** | 9 | **.1** | 2 | **.0** | 44 | **.7** | 8th larva whole body |
| SRR5286154 | 33964933 | 20 | **.6** | 11 | **.3** | 2 | **.1** | 12 | **.4** | 3rd larva whole body |

**Table S1.** Analysis of SIFamide-related gene expression in *Periplaneta* transcriptome SRAs. Number of half-reads in the various transcriptome SRAs available at NCBI. SRA indicates the SRA identifiere and Spots the total number sequences present in each SRA. The number of reads for the four genes are indicated in light blue, their relative number, i.e. their number per million reads are indicated in bold black. The final column gives a short description of the tissues analyzed. Note that the SIFamide gene is strongly expressed in the brain, but also in the subesophageal ganglion and chemosensory organs. The SMYamide gene is strongly expressed in the subesophageal ganglion, very little in the brain, but also in the chemosensory organs. The number of reads for the receptors are much smaller. Apart from the brain, there is a significant number of reads for SIFaR1 in the testes. Note however that the whole female body also has a significant number of reads for both SIFamide receptors and these reads are unlikely to be derived from the brain, as there are relatively few SIFamide reads in that particular SRA (SRR5286151).

```
Nilaparvata-2     ------------------------------------------------------------
Zootermopsis-2    ------------------------------------------------------------
Blattella-2       ------------------------------------------------------------
Periplaneta-2     ------------------------------------------------------------
Ixodes-1          ------------------------------------------------------------
Ixodes-2          ------------------------------------------------------------
Drosophila        -----------------------MAVNGRMRKRKHRSHTAGDMPTTTAPTTTT-------
Locusta           ---------------------------------RRRSGGSARCA-CAGAAAAPGGG-GAA
Nilaparvata-1     -----------------MMAPLRLP-----------------------------------
Bombyx            ----------------MKMAPLRLPVDYYTDDFLNFSTQN---------PNNERHHTRHN
Apis              --------MLQPPSPLLEMASLRLPDSEEFDVDPRRRSNNGNGPATTVSAVPPATVSTLL
Lednia            ------------------------------------------------------------
Rhodnius          ------------------MVSLRLPDSEEYIEARGRALETGLAPTRA-------------
Calopteryx        ------------------------------------------------------------
Isoperla          ------------------------------------------------------------
Blattella-1       MADLVESPAYHASSPLLEMASLRLPDSEEYEFENRRRADSLFTSSSTDSSSNTLHLPTRS
Periplaneta-1     MADPNIPPGYHASSPLLEMASLRLPDSEEYEFENRRRADSLFTSSSTDSSSSTLHLSTRS
Zootermopsis-1    ------------------------------------------------------------
Anisolabis        ------------------------------------------------------------
Medauroidea       ------------------------------------------------------------
Dryococelus       ------------------------------------------------------------
Timema            ------------------------------------------------------------
Tribolium         -------------MGVLDLAALRIDDEEEFYDERRRRA----------------------
Laupala           ------------------------------------------------------------
Teleogryllus      ---MAEAGGLPASSPLLEMASLRLPDSEELDLDARRRGDAAG---VLTSSSTPLHLAARA
```

```
Nilaparvata-2     ------------------------------------------------------------
Zootermopsis-2    ------------------------------------------------------------
Blattella-2       -------------------------------------MTTSI------------------
Periplaneta-2     ------------------------------------------------------------
Ixodes-1          ----------------------------MRRAN-LTFNCYGLYHWTLSTL----------
Ixodes-2          ----------------------------MMRTRPPAMLGSSLLNADNVT-----------
Drosophila        ---------TAGNGSDSGSFSSTSSAIKLSNGAITDTLLAAVLTTATATVAPAASSLISS
Locusta           EERRQCALRVRGGGGGSGRWRSG------GG--AAAVPGAPPLAMDA-------------
Nilaparvata-1     ------------------------------------------------------------
Bombyx            ----------------HSHLRES------HKNHVADMLSNSII--DA-------------
Apis              ----------------------Q------MRNEVGDYLNSLIV--EA-------------
Lednia            ------------------------------------------------------------
Rhodnius          ----------SN--TSSRRLFV--------NSLLVDFVMDTLV--TN-------------
Calopteryx        ------------------------------------------------------------
Isoperla          ------------------------------------------------------------
Blattella-1       QPYRQLPWNIS-----NRRILSE------SGTKLVDFFGNTVL--EL-------------
Periplaneta-1     QAYRQLPWNVS-----NRRI-SE------GGNKLVDFIGNTYL--DL-------------
Zootermopsis-1    -------------------M-S------------ETFENATCV-----------------
Anisolabis        ------------------------------------------------------------
Medauroidea       ----------------------------------------------M-------------
Dryococelus       ----------------------------------------------M-------------
Timema            ------------------------------------------------------------
Tribolium         HLLAAVPMPEPNE-TM----------------LTAEYLGNLFD--VV-------------
Laupala           ------------------------------------------------------------
Teleogryllus      HAYRPAPAPHPLS-NFSRSIVAE------SSGYLTDFVGNTIK--VA-------------
```

Fig. 3. Sequence alignment of various SIFamide GPCR homologs.

```
Nilaparvata-2    ----------------------------------------MFLPLSV---PRTQVI----
Zootermopsis-2   --------------------------MDTSNECHPNRSFCDWWPNMT---SAEV------
Blattella-2      ------------------------INMDITDECRPNRSCPSWWTNAT---VADNDS----
Periplaneta-2    --------------------------------CRPNQSRCNWWPNMT---SEYHEL----
Ixodes-1         -----------------VPALTGTFTSGAGKSSDS--GNGSAFPMLATA----AVPLSVT
Ixodes-2         ------------------------LRWGDSQTDDT--SQSET-------------ANAVQ
Drosophila       MSAAAATTTATSSQLAVVSTMQA-VALPGVSIPDA--TSSTYYANLLSMSPATTSLISVA
Locusta          -SGSGSGPSAGGAVAVAVAVRHN-ASAGGATAAAG--GATGA-------------VA--A
Nilaparvata-1    ------------------------------LAEEG--SAAAF-------------GL--N
Bombyx           -FNTRFVEN---SVL---PDM------------EP--LSSHM-------------DL--E
Apis             -IATSPG-----SR--------G-PGM----DEHE--GASSA-------------LL--N
Lednia           ------------------------------------------------------------
Rhodnius         -TTPSASSS----------------------PSSP-------------------------
Calopteryx       ------------------------------------------------------------
Isoperla         ------------------------------------------------------------
Blattella-1      -LTMNNNHHHNGSAT---PSLHE-NGMDSIILEDS--GNSS--------------SI--N
Periplaneta-1    -VTMANNNNNNISF----NAFHD-GVMEASLQDES--AS----------------GF--N
Zootermopsis-1   -LFSAMTNSKNGSL----NAFHG-SGIDGSVHDEP--GS----------------GL--N
Anisolabis       ------------------------------------------------------------
Medauroidea      -ALAGFN---------------N-LTVDLLNHSRA--DNGT--------------AW--L
Dryococelus      -AVAALNASA------------D-FFLGVGNLSRP--GNGT--------------PW--G
Timema           ---------M------------N-NGSGGGLVNED--AGGF--------------LW--E
Tribolium        -AHTESTK----------------------------------------------------
Laupala          ---------------------------------KG--DGAD--------------GI--F
Teleogryllus     -TNHSAAG------------------HDA--ADKG--EQGD--------------AL--Y
```

```
Nilaparvata-2    ----Y-----TAS--TGATGEPSRNELRYSVVLTVVFCVAYLLVFCVGVVGNFAVVAVVC
Zootermopsis-2   ----Y-----NVSRIGPTTVTYVVYKFRHPLSVTVFFCVAYALVFAVGVVGNCFVVGVVY
Blattella-2      ----F-----NVS----RNHTFIVYEFRHSLTVTISFCIAYALVFVIGVVGNCFVVGVVY
Periplaneta-2    ----L-----NGS-LNGSTPTFAVFELRHSLSVTISFCIAYAVVFVVGVVGNCFVVGVVY
Ixodes-1         Q---------EPPDTLVSEDWVDACFLRHTAPWAVAYCVAYSVVFVVGIVGNCCVLKVVS
Ixodes-2         DNGSW------NASDYDLYSIPSDLWMRYSPGIVAVFCLAYSVVFVMGLLGNSFVVAVVA
Drosophila       ATKSYNDSVLRWEQLDGNGDFGFDPLYRHSLAMSIAFCVAYILVFLVGLIGNSFVIAVVL
Locusta          SA-NASSA-VMAAAAGPWHLLPDDLVYRHSAAMTAVYCAAYSLVFLLGLVGNCCVIAVVW
Nilaparvata-1    VT-RRLR--MDPAASNMSEPFWHDALYRHSLGMSVVFCAAYLIVFVVGLVGNVFVIAVVY
Bombyx           ER-HYPS--RMNGTLNRSDFGGDEFMYRHSGAMTAVYCAAYLLVFLVGLVGNCFVIAVVY
Apis             AS-KTAAA-AANLTAGDGQIPAVDRLYRHSMAMSAVYCVAYVLVFVVGLIGNSFVIAVVY
Lednia           -----------SNLNNVTSDIFPEYFYRHSPVLTVTFCIAYLIVFVVGMCGNSMVINVVF
Rhodnius         ----AAGA-VADLPDSTSNQTYQHLFYRHSIAMTIVFCVAYLIVFIVGLIGNCFVIMVVY
Calopteryx       ---ALANS-SNATASLLAERLPTELFYRHSLAMTVVYCVAYLIVFAVGLVGNCFVIAVVY
Isoperla         -------------NNSTNSTYMPDLFYRHSLAMTVVYCIAYLIVFAVGLVGNCFVIAVVY
Blattella-1      S--SLVN--G-TLATGMNFSMDPIPFYRHSLPMTAVYCFAYILVFAVGLVGNCFVIAVVY
Periplaneta-1    A--SNVN--A-TL----GNATEPIPFYRHTFAMTAVYCFAYILVFAVGLVGNCFVIAVVY
Zootermopsis-1   E--SILN--A-TA----ANGTDPIPFYRHSFAMTAMYCFAYILVFAVGLVGNYFVIAVVY
Anisolabis       -------N-V-T--GDSLNATVVTEFYRHSVAMTAIYCVAYLIVFAVGLVGNCLVIAVVY
Medauroidea      DD-GYGG-------------LPTEMLYRHGFAMTVVYCVAYLIVFAVGLVGNSFVVAVVY
Dryococelus      DD------------AGNATFAFDELVYRHSFAMTAVYCVAYLLVFAVGLVGNCFVIAVVY
Timema           DY-GGYNL-T-YGNGTNGSFSYDEMVYRHTFAMTAIYCVAYLLVFAVGLVGNCFVIAVVY
Tribolium        ----WAN----ATLNATETAFVPELFYRHSMAMTAVYCVAYLLVFAVGIVGNFFVIAVVF
Laupala          NG-TFGI----NATTNGTMTLSGDLFYRHTLVMTAVYCVAYVLVFAVGLIGNCFVIAVVF
Teleogryllus     NG-SLAL----NASGNVTAALGADLFYRHSLAMTAVYCVAYVLVFAVGLIGNCFVIAVVF
```

Fig. 3. Sequence alignment of various SIFamide GPCR homologs.

```
Nilaparvata-2   RSPRMRTP-TNLFIANLACADLLVNIICLPFTLISNIMTAWTMGWLVCKTIPYMQGVSVN
Zootermopsis-2  RSPRMRSP-TNLFIANLACADLLVNVLCLPFTLVGNIMSAWIMGWVICKTVPYLQGVSVS
Blattella-2     RSPRMRSP-TNLFIANLACADLLVNVLCLPFTLVGNIMSAWIMGWVICKTVPYLQGVSVS
Periplaneta-2   RSPRMRSP-TNLFIANLACADLLVNVLCLPFTLVGNIMSAWIMGWVICKTVPYLQGVSVS
Ixodes-1        RNTSMKNSAFHMLLVNLAVADLMVIVFCLPVTLVGHLFGPWILGLFVCKGVSYLQGVSVC
Ixodes-2        RSPRMRTV-TNYFIVNLAMADILVVVFCIPATLVSNIFVPWVLGWFMCKTMSYLQGVAVS
Drosophila      RAPRMRTV-TNYFIVNLAIADILVIVFCLPATLIGNIFVPWMLGWLMCKFVPYIQGVSVA
Locusta         RSPRMRTV-TNLFIANLAAADLLVVVVCLPATLVSNIFVPWLLGWWMCKTVPYVQGVSVA
Nilaparvata-1   RSPRMRTV-TNFFIVNLAIADILVVVFCLPATLLSNIFVPWILGWWMCKTVPYVQGVSVA
Bombyx          RSPRMRTV-TNFFIVNLAFADILVIVFCLPATLMSNIFVPWVLGWLMCKTVPYVQGLSVA
Apis            RSPRMRTV-TNFFIVNLAVADVLVIVFCLPATLLSNIFVPWLLGWFMCKAVAYIQGVSVA
Lednia          RSPRMWTV-TNYFIVNLAFADILVIVFCVPATLMSNIFVPWILGWWMCKMVPYVQGVSVA
Rhodnius        RSPRMRNV-TNFFIVNLAVADILVIVFCLPATLMSNIFVPWVLGWWMCKTVPYVQGVSVA
Calopteryx      RSPRMRTV-TNYFIVNLAAADILVIVFCLPATLMSNIFVPWVLGWWMCKTVPYVQGVSVA
Isoperla        RSPRMRTV-TNFFIVNLAIADILVVAFCIPATLTANIFVPWVLGWFMCKTVPYIQGVSVA
Blattella-1     RSPRMRTV-TNFFIVNLAVADILVIVFCLPATLLSNIFVPWLLGWWMCKTVPYVQGVSVA
Periplaneta-1   RSPRMRTV-TNFFIVNLAVADILVIVFCLPATLLSNIFVPWLLGWWMCKTVPYVQGVSVA
Zootermopsis-1  RSPRMRTV-TNFFIVNLAVADILVIIFCLPATLLSNIFVPWLLGWWMCKTVPYVQGVSVA
Anisolabis      RSPRMRTV-TNFFIVNLAVADILVIVF-LPATLVANIFVPWILGWWLCKTIPYVQGVSVG
Medauroidea     RSPRMRTV-TNYFIVNLAVADILVIVFCLPATLMANIFVPWVLGKWMCKTVPYVQGVSVA
Dryococelus     RSPRMRTV-TNYFIVNLAVADILVIVFCLPATLMANIFVPWVLGWWMCKTVPYVQGVSVA
Timema          RSPRMRTV-TNFFIVNLAVADILVIVFCLPATLMANIFVPWVLGWWMCKTVPYVQGVSVA
Tribolium       RSPRMRTV-TNFFIVNLAVADILVIVFCLPATLMSNIFVPWVLGWWMCKTVPYIQGVSVA
Laupala         RTPRMRTV-TNFFIVNLAVADILVIVFCLPATLMSNIFVPWVLGWWMCKTVPYVQGVSVA
Teleogryllus    RTPRMRTV-TNFFIVNLAVADILVIVFCLPATLMSNIFVPWVLGWWMCKTVPYVQGVSVA
```

```
Nilaparvata-2   ASINTLVAISFERWLAICYPMRWQMTSRVCKLVILLIWLFSLTITLPWALFFQLRPMGD-
Zootermopsis-2  ASINTLVAISVERCLAICYPLKWQMTSRACRVVVIIIWTFSLTITLPWAIFFGLKPLEE-
Blattella-2     ASINTLVAISVERCLAICYPLKWQMTSRACRVVVVIIWTFSLIITLPWAIFFGLRPLEE-
Periplaneta-2   ASINTLVAISVERCLAICYPLKWQMTSRACRVVVVIIWTFSLIITLPWAIFFGLHPLED-
Ixodes-1        ASVNTLVAISIDRFFAICHPMKRQITIRVCRVIIAVIWSFSLTITLPWTFFFRLMPMLSE
Ixodes-2        ASINTLVAISMDRCLAICYPLKCQLSTRSVRKILVIIWTFSIAITFPWALYFTLQPLHPS
Drosophila      ASVYSLIAVSLDRFIAIWWPLK-QMTKRRARIMIIGIWVIALVTTIPWLLFFDLVPAEEV
Locusta         ASVYSLIAVSLDRFLAIWWPLKFQITKRRARLIIAAIWLLALTTTLPWAVFFDLVSVLD-
Nilaparvata-1   ASVFSLIAVSIDRFLAIWYPLKCQITTRRARTIIAVIWLVAATTTLPWALFFDVV-IFSE
Bombyx          ASVYSLVAVSLDRFLAIWWPLKCQITKRRSRMMIVFIWIFAILVTTPWVFFFDLVVVFEE
Apis            ASVYSLVAVSLDRFLAIWWPLKCQITKRRARMIIVVIWFIALTTTSPWLLFFDLVAIYKD
Lednia          ASVYSLIGVSLDRFLAIWWPMKCQITKRRARFIIFLIWVIAFATTIPWAIYFDLVVM--E
Rhodnius        ASVYSLIAVTLDRFLAIWWPLKCQITTRRARLMILVIWVVALTTTIPWALFFDLVVIFTD
Calopteryx      ASVYSLIAVTLDRFLAIWYPLKCQITKRRARLMIFIIWVVALTTTIPWALFFDLVIIFRE
Isoperla        ASVYSLIAVSLDRLLAIWWPMKCQITKRRARIMIFVIWVIALSTTIPWAIYFDLV-IFHE
Blattella-1     ASVYSLIAVSLDRFLAIWWPLKCQITKRRARFMIFVIWVIALTTTIPWALFFDLVIIFND
Periplaneta-1   ASVYSLIAVSLDRFLAIWWPLKCQITKRRARFMIFVIWVIALTTTIPWALFFDLVMVFND
Zootermopsis-1  ASVYSLIAVSLDRFLAIWWPLKCQITKRRARFMIFFIWVIALTTTIPWALFFDLVIIFND
Anisolabis      ASVYSLIAVSVDRFLAIWWPLKCQITKRRARVMIVIIWVIALTTTIPWALFF----VFRD
Medauroidea     ASVYSLIAVSADRFLAIWWPLKCQITKPRARFVIVLIWVIALSTTIPWALYFELVSVFND
Dryococelus     ASVYSLIAVSLDRFLAIWWPLKCQITKRRARFMIVLIWVIALTTTIPWAVFFDLVSIFRD
Timema          ASVYSLIAVSLDRFLAIWWPLKCQITKRRARIMIVFIWVIALTTTIPWAVFFDLVSIFRD
Tribolium       ASVYSLIAVSLDRFLAIWWPLKCQITKKRARLMIVIIWFIALTTTIPWAIFFDLVAVFND
Laupala         ASVYSLIAVSLDRFLAIWWPLKCQITKRRARFIILVIWVIALTTTIPWALFFDLVQIYAD
Teleogryllus    ASVYSLIAVSLDRFLAIWWPLKCQITKRRARFIIFVIWVIALTTTIPWALFFDLVVIYSE
```

Fig. 3. Sequence alignment of various SIFamide GPCR homologs.

```
Nilaparvata-2     ----------GSSMQTCLETWPTPYSE-NVYFVLANLVMCYLLPLALISICYFRIWRRVCC
Zootermopsis-2    ----------GSDVLICTESWPSPHSG-NIYFVVAHLVMCYLFPLTLISLCYLLIWRRVCR
Blattella-2       ----------GSELQICTESWPSPDSG-NVYFVVAHLVMCYLFPLTLISVCYLLIWRRVCR
Periplaneta-2     ----------GSDLQICTESWPSQDSG-NIYFVVAHLVMCYLFPLTLISVCYLLIWRRVCR
Ixodes-1          ---------SNSSLQVCREDWPTERMG-MLYFIVANLILCYLLPLCVITLCYIFIWLKVWR
Ixodes-2          ----------IPGISLCVEQWPDETSS-TLYFILAHLVLCYLFPLLLIIVCYSCIWVKVWR
Drosophila        FSDALVSTYTQPQYLCQEVWPPGTDG-NLYFLLANLVACYLLPMSLITLCYVLIWIKVST
Locusta           ------------DVRVCVEVWPDPTDG-ALYFLFGNLLCFYVVPFALISLCYVLIWIKVCK
Nilaparvata-1     ----------EPDVPMCVEVWPDYLNG-NLYFLFANLIMCYIVPMILISLCYILIWIKVCK
Bombyx            ----------NPNVHLCIDVWPNPLSE-VLYFVVGNLIFCYILPMVMITMCYILIWIKVWR
Apis              ----------DPDLRLCLEVWPRPKDE-TLFFLIGNLTLCYVLPTILISLCYILIWIKVWR
Lednia            ----------ELDVSLCIEVLPDPITG-ALNFLIANVLLCYILPTILISMCYVLIWIKVSK
Rhodnius          ----------NPEVKVCSEVWPEYLNG-SLYFLIANLLFCYILPMILISMCYVLIWIKVCK
Calopteryx        ----------MPETRLCLEVWPPSLNG-DLYFLLANLVFCYVLPTILICLCYILIWIRVAR
Isoperla          ----------VPDVKLCMEVWPDSTNGGVLYFLIVNLLLFYILPMILISMCYVLIWVKVWK
Blattella-1       ----------APDIQLCLEVWPDSLDG-ALYFLIANLLFCYILPMILISLCYILIWVKVWK
Periplaneta-1     ----------APDVLLCVEVWPDALDG-ALYFLIANLLFCYILPMVLISLCYILIWVKVWK
Zootermopsis-1    ----------APDVQLCVEVWPDALDG-TLYFLIANLLFCYILPMILISLCYILIWVKVWK
Anisolabis        ----------LPDNQLCQEVWPDPMDG-ALYFLIANLMFFYILPMILISLCYILIWIKVWR
Medauroidea       ----------APDVQLCIEIWPENMDG-NLYFLIDNLIICYILPLIVISICYILIWIKVWR
Dryococelus       ----------APEIQLCLEVWPDSMDG-DLYFLIANLVFCYILPMILISLCYILIWIKVWR
Timema            ----------APEIQLCLEVWPEALDG-DLYFLIANLVFCYILPMILISMCYVLIYIKVWR
Tribolium         ----------APDVQLCIEVWPANMDG-ALYFLIANMVFCYILPMILITMCYVLIWIKVWR
Laupala           ----------APEVRLCVEVWPDSMNG-ALYFLIANLMFCYLLPMVLISLCYILIWIKVWR
Teleogryllus      ----------APDVLLCVEVWPDSMNG-ALYFLIANLMFCYLLPMVLISLCYILIWIKVWK

Nilaparvata-2     RKMPGEVQM---YQELIIHRSKVKVIKMLFIVIVLFACSWLPLYVIFTRLKLGGDLQP--
Zootermopsis-2    RTLPGEPHSHGGVVDLMIHRSKVKVIKMLLVVVISFALSWLPLYVLFTRVKFGGPFSS-E
Blattella-2       RTLPGEPHPQGGVMDMMIQRSKVKVIKMLLVVVVSFALSWLPLYVLFTRVKFGGPFSE--
Periplaneta-2     RTLPGEPHPPGGVMDVMIHRSKVKVIKMLLVVVVSFALSWLPLYVLFTRVKFGGPFSS-E
Ixodes-1          RRPPGE-AH-DLGVENMIQRSKVKVAKMLLVVVIVFAISWLPLYAIFARLKIGEPLVDGS
Ixodes-2          RSIPGE----SKHTEIMVQKSKLKVVKMMFVVVVIFVISWLPLYIIFTRIKLDSPPEEGS
Drosophila        RSIPGEMSK-DAQMDRMQQKSKVKVIKMLVAVVILFVLSWLPLYVIFARIKFGSDISQ--
Locusta           RDIPGD-ST-DAQLERMQHKSKIKVVKMLVVVVILFLASWLPLYVIFARIKLGGELAA--
Nilaparvata-1     RHIPSD-TK-DAQMERMQQKSKVKVVKMLVAVVILFVLSWLPLYAIFTRIKLGGDLDV--
Bombyx            RSIPTD-TQ-DAQMERMQQKSKVKVVKMLVAVVILFVLSWFPLYLIFARIKLGGPIKK--
Apis              RHIPSD-TK-DAQMERIQQKSKVKVVKMLVVVVILFVLSWLPLYVIFTVIKLGDEQ----
Lednia            RDIPTD-TK-DAQIERMQQKSKVKVVKMLVVVVILFVLSWLPLYLIFASIKLGWDLSR--
Rhodnius          RHIPSD-SK-DAQMERMQQKSKVKVVKMLVVVVILFVLSWLPLYLIFARIKLGGEISG--
Calopteryx        RHIPTD-SK-DAIAERMQQQSKVKVIKMLVAVVVIFVLSWLPLYAIFARIKLGGEVED--
Isoperla          RHIPSD-TK-DAQIERMQQKSKVKVVKMLVVVVILFVLSWLPLYIIFARIKLGGDIER--
Blattella-1       RTIPTD-TK-DAQMERMQQKSKVKVVKMLVAVVILFVLSWLPLYVIFARIKLGGEVEI--
Periplaneta-1     RTIPTD-TK-DAQMERLQQKSKVKVVKMLVAVVILFVLSWLPLYVIFARIKLGGDVEI--
Zootermopsis-1    RTIPTD-TK-DAQMERMQQKSKVKVVKMLVAVVILFVLSWLPLYVIFARIKLGGDVEM--
Anisolabis        RDIPTD-TK-DAQMERLQQKSKIKVVKMLVIVVILFVLSWLPLYVICARIKLGGEREL--
Medauroidea       RDIPTD-TK-DAQMERLQQKSKVKVVKMLVAVVILFVLSWLPLYVIFTRIKLGERISR--
Dryococelus       RDIPTD-TK-DAQMERLQQKSKVKVVKMLVAVVILFVLSWLPLYVIFARIKLGGRTSR--
Timema            RDIPTD-TK-DAQMERMQQKSKVKVVKMLVAVVILFVLSWLPLYTIFARIKLGGKTAR--
Tribolium         RHIPTD-TK-DAQMERMQQKSKVKVVKMLVAVVILFILSWLPLYVIFARIKFGGHIEA--
Laupala           RDIPTD-TK-DAQMERMQQKSKVKVVKMLVAVVILFVLSWLPLYVIFARIKLGGETEL--
Teleogryllus      RDIPTD-TK-DAQMERMQQKSKVKVVKMLVAVVILFVLSWLPLYVIFARIKLGGETEL--
```

Fig. 3. Sequence alignment of various SIFamide GPCR homologs.

```
Nilaparvata-2     WEEPLVYNLLPLAQWLGASNSCINPVLYAFFNKKFRAGFKAILSSKSCFTTLRYDTYSFD
Zootermopsis-2    TEEATIHSLLPVAQWLGASNSCVNPILYAFFNRKFRIGFKEIITSRSCCSTLRYNNEFR-
Blattella-2       SEEAAVHAILPVAQWLGASNSCVNPILYAFFNKKFRAGFKAIVTSKSCCTPIRYNNEYYS
Periplaneta-2     TEEAAIHALLPIAQWLGASNSCVNPLLYAFFNRKFRIGFKAILTSRSCCKPLRYNNDFS-
Ixodes-1          AEQAVIEVAAPVAQWLGASNSCINPILYAFFNSKFRMGFKAILLKCFCPKKFSSRQERSC
Ixodes-2          VEWNLMLILTPVAQWLGASNSCINPVLYAYFNQKFRKGFLAIIKSRSCCGTLREPSY--S
Drosophila        EEFEILKKVMPVAQWLGSSNSCINPILYS-VNKKYRRGFAAIIKSRSCCGRLRYYDN--V
Locusta           WEDEALPVATPVAQWLGASNSCINPLLYAFFNHKFRRGFAAILRSRRCCGRLRYYET--V
Nilaparvata-1     WETKLFVVLTPVAQWLGSSNSCINPILYAFFNNKYRRGFSAILRSRQCCGTLRYYDT--V
Bombyx            WEEEMLPIVTPLAQWLGASNSCINPILYAFFNKKYRKGFVAIIKSRKCCGRLRYYET--I
Apis              REDEIVPIATPIAQWLGASNSCINPILYAFFNKKYRRGFVAILKSGRCCGKIRYYET--V
Lednia            GEEDFLQYATPIAQWLGSSNSCINPILYAFFNKKFRRGFVAIWKSKRLCGRLRYYET--V
Rhodnius          WEEDMLPMATPVAQWLGASNSCINPILYAFFNNKYRRGFAAILKSRKCCGTLRYYDT--V
Calopteryx        WEDKILPVATPLAQWLGASNSCINPVLYAFFNKKYRRGFVAIVKSRKCCGRLRYETA--I
Isoperla          WEEEILPIATPIAQWLGASNSCINPILYAFFNKKYRRGFVAILKSRRCCG-LRYYET--I
Blattella-1       WEDDILLVATPIAQWLGASNSCINPILYAFFNKKYRKGFIAILKSRRCCGRLRYYES--V
Periplaneta-1     WEDDILLVATPIAQWLGASNSCINPILYAFFNKKYRKGFIAILKSRRCCGRLRYYES--V
Zootermopsis-1    WEDDILLVATPIAQWLGASNSCINPILYAFFNKKYRKGFIAILKSRRCCGHLRYYES--V
Anisolabis        WEEDILTFVFPIAQWLGASNSCINPILYAFFNKKYRRGFIAILKSRKCCGRLHYYES--V
Medauroidea       SEEEFIQVVTPIAQWLGSSNSCINPILYAFFNKKYRRGFVAILKSRRCCGRLRYYES--V
Dryococelus       WEEELIQVVTPIAQWLGSSNSCINPILYAFFNKKYRRGFVAILKSRRCCGRLRYYES--V
Timema            WEEDVIQVVTPIAQWLGSSNSCINPILYAFFNKKYRRGFVAIFKSRRCCGRLRYYES--V
Tribolium         WEEEILPIATPIAQWLGASNSCINPILYAFFNKKFRRGFVAIIKSRRCCGRLRYYET--V
Laupala           WEEDILPIATPIAQWLGASNSCINPILYAFFNKKYRRGFIAILKSRKCCGRLRYYET--I
Teleogryllus      WEEDVLPIATPIAQWLGASNSCINPILYAFFNKKYRRGFIAILKSRKCCGRLRYYET--I

Nilaparvata-2     GR------NL-SSNY-GNRNDTVKR--PLTRMTISSSAALRTGRIG--GSEGSLVRTR--
Zootermopsis-2    -A------SV-NSSF-SCRTNTKNGSVLMTRMTITKSKGGKS-VRR--TRAGSLHRHN--
Blattella-2       -T------SV-KSTF-SCRMNTKNGSILMSKKALNPSSSTKT-RVP--TTGSNLQRHH--
Periplaneta-2     ------------SSF-SCRMNGS---------------------VM--APASNLQRHN--
Ixodes-1          KMLSNTRTSM-------------------------TAGRRKDSRMSDVF-----------
Ixodes-2          VRG----TTLRS-----GNT-RG-LSR-SENLEYVEHPRAHSK-AGTLLINGGPDAQA--
Drosophila        AIASS-TTSTRKSSHYHPSSSRK---------SPSSPGLRKTNAVSYIYEHNSLRRHNLM
Locusta           ALSGS---SVRKSF---NNN---NSS-----------TRRPDTSVSYIFNNTGV------
Nilaparvata-1     AMANSSSASVRKSSCYVNSNNPSVRRQ-------IMHQASQDSAVSYISNNTGV------
Bombyx            ALQSS-STSTRKSWHYNNNNP---SIT-------RRSPPVDKNAVSFIFSHTGV------
Apis              AMMSS-STSMRKSSYYVNNNNNNNNNS-STRRTFHGPPVHQESNVSYIFNHTGV------
Lednia            AMMSSSASSTRKSSYYVNNNN--SSTR-------RSPPVAQDSAVSYISNNTGI------
Rhodnius          VRANSSSTSLRKSSYYVTNNNNNNNSS-------TRRQLSQDTNVSYISNNTGV------
Calopteryx        ------TSSLRKSSYYMNS------------------SMRQDTSVAYIYNNTGV------
Isoperla          ALMSSSGSSMRKSSYYVNN----NSST-------RRPPAGQDTSVSYICNNTGI------
Blattella-1       -AMSS-SASLRKSSYYVNNN---NSST-------RRPPPGQDTSVSYIFNNTGV------
Periplaneta-1     -AMSS-SASMRKSSYYVNNN---NSST-------RR-PPGQDTSVSYIFNNTGV------
Zootermopsis-1    -AMSS-TTSMRKSSYYVNNN---NSST-------RR-PPGQDTSVSYIFNNTGV------
Anisolabis        ALMSS-STSLRKSSYYVTNNN--NNST-------RRPPPGQDTSVSYIFNNTGV------
Medauroidea       -MMSS-SASLRKSSYYVNN----NNST-------RRPPPGQDTSVSYIYNNTGV------
Dryococelus       -MMSS-SASLRKSSYYVNN----NNST-------RRPPPGQDTSVSYIFNNTGV------
Timema            -MMSS-SASLRKSSYYVNN----NSST-------RRPAPGQDTSVSYIFNNTGV------
Tribolium         AMMSS-STSMRKSSHFYNN-----NSS-------TRKLPLQDNAVSYIYNNTGV------
Laupala           AMMSS-STSMRKSSYYTNN-----NSS-------TRRPPGQDTSVSYIFNNTGV------
Teleogryllus      AMMSS-STSMRKSSYYTNNN----NSS-------TRRPPGQDTSVSYIFNNTGV------
```

Fig. 3. Sequence alignment of various SIFamide GPCR homologs.

```
Nilaparvata-2     -----TLSFKAMNQVN-NNSSE-GPFYTEKV-------------------NGLTLSDMTL
Zootermopsis-2    -----SASAAAVQRIN-KNNEQFRKSYSRTCRNSISEAGENHRII--KSSSGS---HL--
Blattella-2       -----SASTAAVYKIN-KVNENYNNNYTNACKNPTTETG--HHII--KSSSVSALSHL--
Periplaneta-2     -----SASAAAVLKIN-KNN-----NY-TRCKNP---AS--HHIM--KSSSASALSQL--
Ixodes-1          ------------------------------------------------------------
Ixodes-2          ----TNVKAPLVA-----------------------------------------------
Drosophila        MKQDSNLSQQMLLKQDSHGSRQFLIKQESSCSD----ASGTRRLLCQQDSNGSK---VSL
Locusta           ------------------------------------------------------------
Nilaparvata-1     ------------------------------------------------------------
Bombyx            ------------------------------------------------------------
Apis              ------------------------------------------------------------
Lednia            ------------------------------------------------------------
Rhodnius          ------------------------------------------------------------
Calopteryx        ------------------------------------------------------------
Isoperla          ------------------------------------------------------------
Blattella-1       ------------------------------------------------------------
Periplaneta-1     ------------------------------------------------------------
Zootermopsis-1    ------------------------------------------------------------
Anisolabis        ------------------------------------------------------------
Medauroidea       ------------------------------------------------------------
Dryococelus       ------------------------------------------------------------
Timema            ------------------------------------------------------------
Tribolium         ------------------------------------------------------------
Laupala           ------------------------------------------------------------
Teleogryllus      ------------------------------------------------------------


Nilaparvata-2     SANATFV-----------------------------------------------------
Zootermopsis-2    --NATSV-----------------------------------------------------
Blattella-2       --NATAV-----------------------------------------------------
Periplaneta-2     --NATAV-----------------------------------------------------
Ixodes-1          ------------------------------------------------------------
Ixodes-2          ------------------------------------------------------------
Drosophila        SKQDSIVSYMEARRVAALSAQDRSVDSTATQQDTISIESRRPGALPAAATPPSLVDKRQK
Locusta           ------------------------------------------------------------
Nilaparvata-1     ------------------------------------------------------------
Bombyx            ------------------------------------------------------------
Apis              ------------------------------------------------------------
Lednia            ------------------------------------------------------------
Rhodnius          ------------------------------------------------------------
Calopteryx        ------------------------------------------------------------
Isoperla          ------------------------------------------------------------
Blattella-1       ------------------------------------------------------------
Periplaneta-1     ------------------------------------------------------------
Zootermopsis-1    ------------------------------------------------------------
Anisolabis        ------------------------------------------------------------
Medauroidea       ------------------------------------------------------------
Dryococelus       ------------------------------------------------------------
Timema            ------------------------------------------------------------
Tribolium         ------------------------------------------------------------
Laupala           ------------------------------------------------------------
Teleogryllus      ------------------------------------------------------------
```

Fig. 3. Sequence alignment of various SIFamide GPCR homologs.

```
Nilaparvata-2   ------------------------------------------------------------
Zootermopsis-2  ------------------------------------------------------------
Blattella-2     ------------------------------------------------------------
Periplaneta-2   ------------------------------------------------------------
Ixodes-1        ------------------------------------------------------------
Ixodes-2        ------------------------------------------------------------
Drosophila      FVKQDSVISFVDQRPEPRRHQLVKQDSVISFADQRRGLLHKQDSLKTNRSNDAPTHHVSI
Locusta         ------------------------------------------------------------
Nilaparvata-1   ------------------------------------------------------------
Bombyx          ------------------------------------------------------------
Apis            ------------------------------------------------------------
Lednia          ------------------------------------------------------------
Rhodnius        ------------------------------------------------------------
Calopteryx      ------------------------------------------------------------
Isoperla        ------------------------------------------------------------
Blattella-1     ------------------------------------------------------------
Periplaneta-1   ------------------------------------------------------------
Zootermopsis-1  ------------------------------------------------------------
Anisolabis      ------------------------------------------------------------
Medauroidea     ------------------------------------------------------------
Dryococelus     ------------------------------------------------------------
Timema          ------------------------------------------------------------
Tribolium       ------------------------------------------------------------
Laupala         ------------------------------------------------------------
Teleogryllus    ------------------------------------------------------------


Nilaparvata-2   -----------------------
Zootermopsis-2  -----------------------
Blattella-2     -----------------------
Periplaneta-2   -----------------------
Ixodes-1        -----------------------
Ixodes-2        -----------------------
Drosophila      LKKTDSQLSYGTSSPRRNVELYE
Locusta         -----------------------
Nilaparvata-1   -----------------------
Bombyx          -----------------------
Apis            -----------------------
Lednia          -----------------------
Rhodnius        -----------------------
Calopteryx      -----------------------
Isoperla        -----------------------
Blattella-1     -----------------------
Periplaneta-1   -----------------------
Zootermopsis-1  -----------------------
Anisolabis      -----------------------
Medauroidea     -----------------------
Dryococelus     -----------------------
Timema          -----------------------
Tribolium       -----------------------
Laupala         -----------------------
Teleogryllus    -----------------------
```

Fig. 3. Sequence alignment of various SIFamide GPCR homologs.

Fig. S4. Sanger sequence analyis of the RT-PCR product for *Periplaneta* SIFamide receptor 1.

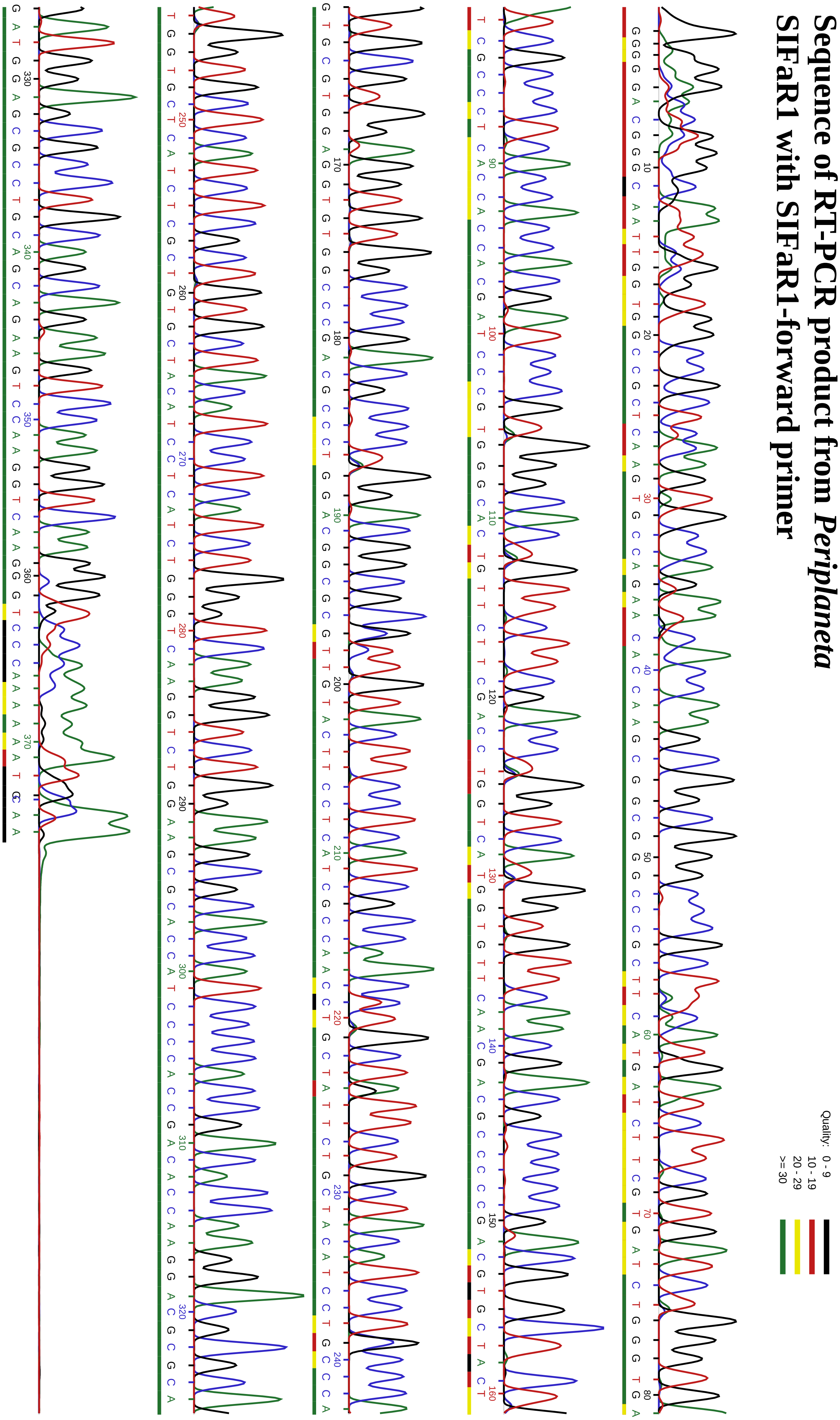

Fig. S5. Sanger sequence analyis of the RT-PCR product for *Periplaneta* SIFamide receptor 2.

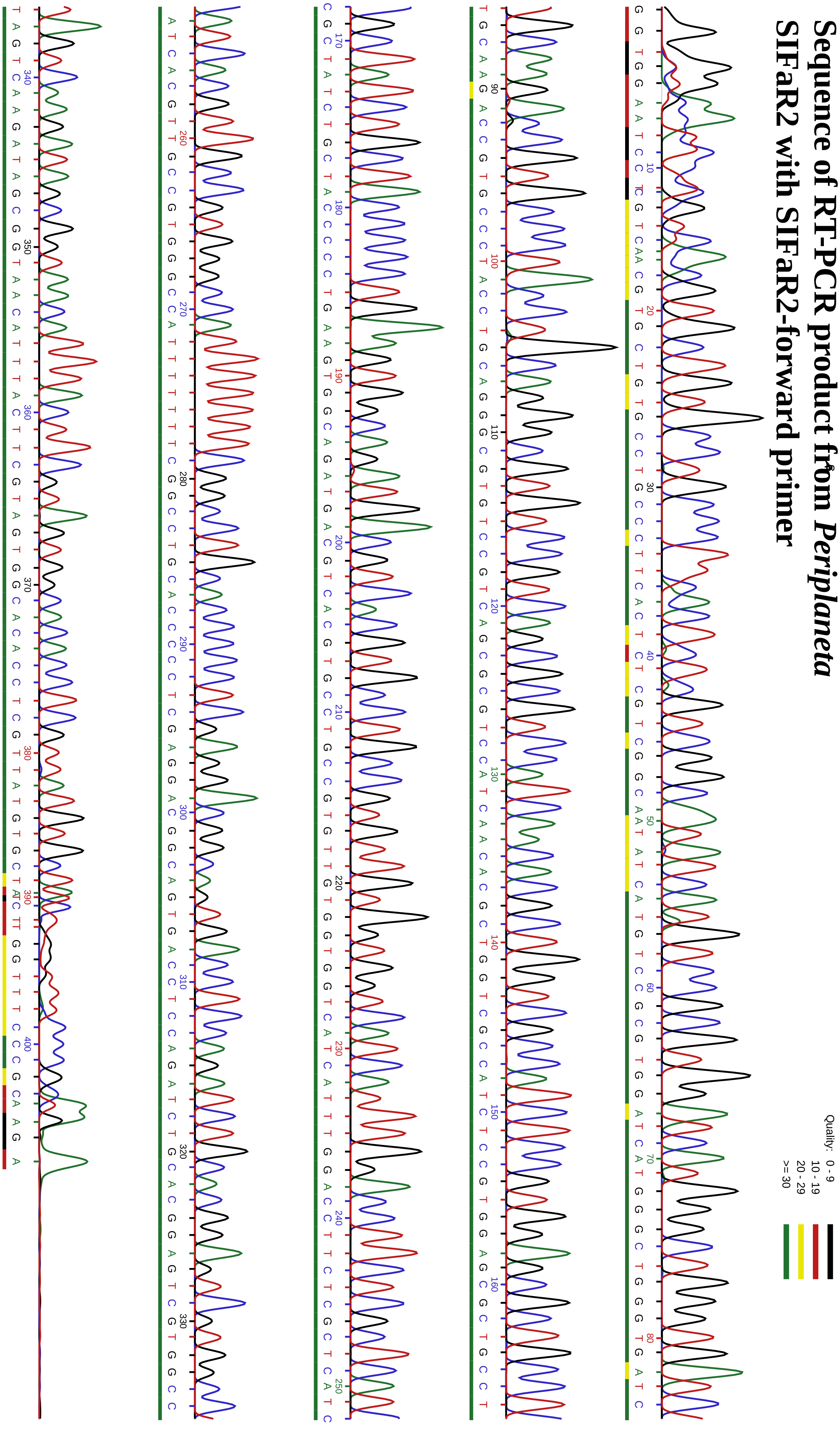
